# GTP-bound *E. coli* FtsZ filaments are composed of Tense monomers: A DNP NMR study using interface detection

**DOI:** 10.1101/2022.04.01.486622

**Authors:** Kelsey M. McCoy, Ann E. McDermott

## Abstract

FtsZ filaments are the major structural component of the bacterial Z-ring and are drivers of bacterial division. While crystal structures for FtsZ from some gram positive bacteria in the presence of GTP-analog like compounds suggest the possibility of a high energy “Tense” conformation, to date it remains an important question to elucidate whether this Tense form is the dominant form in filaments. Using dynamic nuclear polarization (DNP) solid-state NMR and differential isotopic labelling, we directly detect residues located at the inter-monomer interface of GTP-bound WT *Escherichia coli* FtsZ filaments. We combine chemical shift prediction, homology modelling, and heteronuclear dipolar recoupling techniques to characterize the *E. coli* FtsZ filament interface and demonstrate that the monomers in active filaments assume a Tense conformation.

## Introduction

Bacterial cytokinesis and septation are driven by a large set of factors known as the divisome that assemble into the Z ring at the site of division (*1*)(*2*)(*3*)(*4*)(*5*). The major structural component of the Z ring is FtsZ (*6*), a highly conserved member of the tubulin family (*7*)(*8*)(*9*)(*10*) that is present in nearly all known bacterial and archaea genomes (*11*). FtsZ is present in the cytoplasm as a mix of soluble monomers, dimers, and transient oligomers (*12*)(*13*)(*14*), which then assemble into dynamic (“active”) filaments (*15*)(*16*)(*17*) that treadmill around the site of division (*18*)(*19*)(*20*)(*21*), serving as a scaffold for downstream divisome factors (*22*)(*23*), and playing a role in membrane constriction (*24*)(*25*)(*26*).

*In vitro*, purified FtsZ reversibly assembles in the presence of MgCl_2_, KCl, and GTP at pHs between 6.5–7.5 (*27*)(*28*)(*13*). FtsZ filaments assemble cooperatively (*29*)(*18*) with a GTP bound at each monomer interface (*13*)(*30*); the GTPase active site is split such that individual monomers can bind, but not hydrolyze, GTP. Upon GTP binding, the monomers assemble into a straight filaments that are bundled together by a variety of factors. As GTP is depleted, the filaments progressively curve and eventually disassemble. Because FtsZ assembles into single protofilaments composed of monomers, this higher order conformational changes must be due to a corresponding conformational change within the monomers. However, until recently, only a single monomer form was observed using X-ray crystallography.

In *Staphylococcus aureus* FtsZ, a second monomer conformation has been observed either when FtsZ has been bound to a small molecule inhibitor—PC190273 (*31*)(*32*) or after the mutation of specific sites (*18*) including the truncation of the catalytic T7 loop (*33*), and recent evidence suggests that this second “tense” or “T” state can also be trapped in SaFtsZ using GTP mimetics (*34*). In methicillin-resistant *S. auresus*, crystals of wild-type FtsZ contained both T and R states, suggesting an equilibrium between the two states (*35*). Additionally, this second state has been inferred in *Bacillus subtilis* FtsZ using PC190273 bound to a fluorophore (*36*) and it also seems to align with an ~6 Å cryo-EM density map of *E. coli* FtsZ filaments (*18*). This tense or “open” form of the monomer—so call because of the accessibility of the interdomain cleft (*18*)(as shown in figure 1)—is an appealing solution to the two state problem because it maps well to a straight filament (figure 1B). The current model posits that FtsZ monomers preferentially adopt a “relaxed” or “closed” state (the canonical FtsZ structure present in the majority of crystal structures) in the cytoplasmic pool where they form transient dimers, trimers, and oligomers. The tense form then, is the form that is present in active, GTP-bound filaments. As the filaments hydrolyze GTP to GDP, the constituent monomers revert to the relaxed form and the filaments curve and then disassemble. This model allows for the seeding of the cooperatively assembled, treadmilling filaments that are observed *in vitro*.

**Figure 1:**
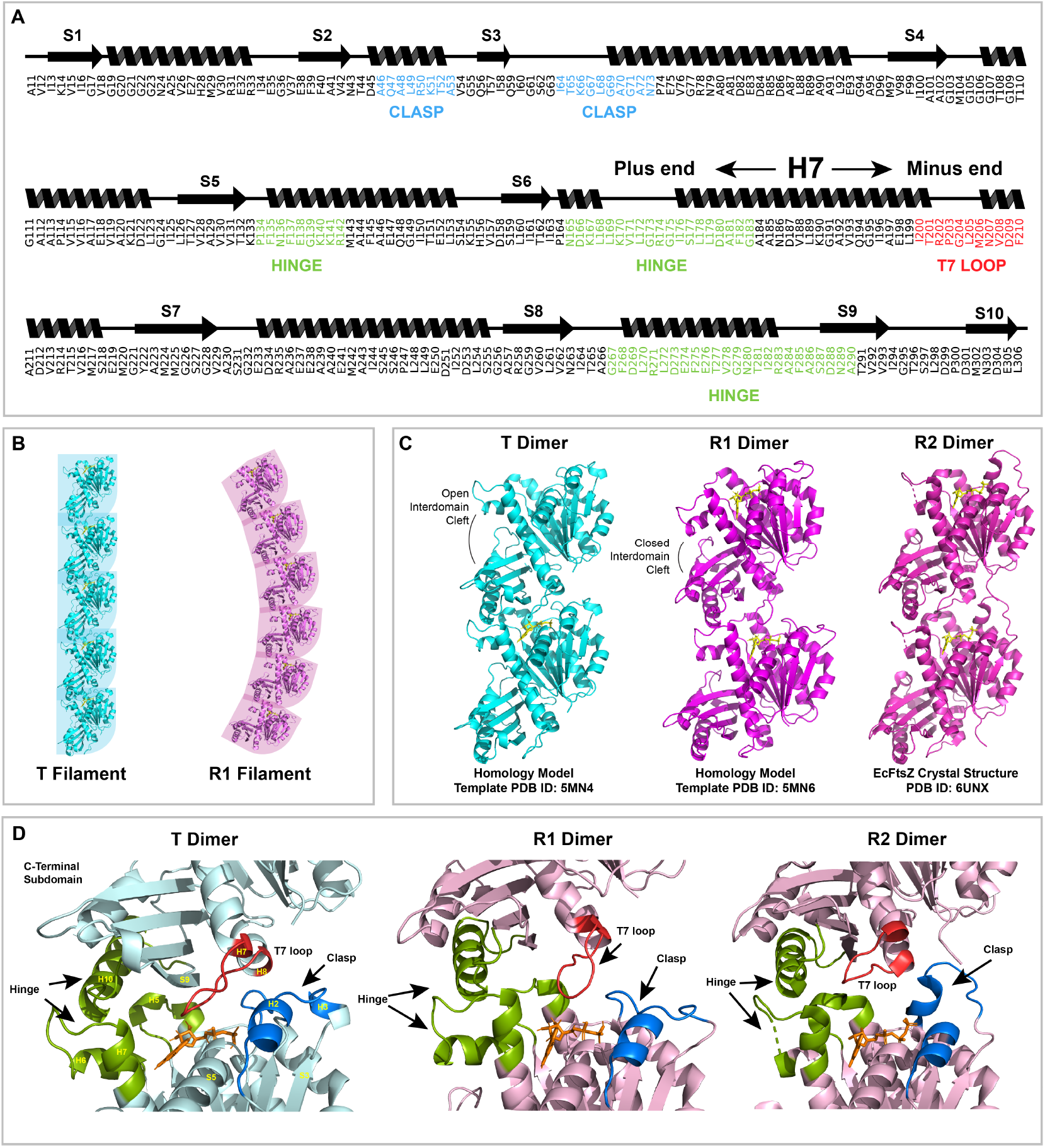
A) Wire diagram and primary sequence of the structured region of E. coli FtsZ used in this study with the interface regions labelled. B) Structural representation of the filament structures of straight and curved filaments as composed of tense and relaxed monomers respectively. PDB ID: 5MN4 (tense monomer) and 5MN6 (relaxed monomer) (*18*). C) Structures of the three E. coli dimer models used in this study. Models T and R1 were constructed by templating the E. coli FtsZ sequence against dimers inferred from the crystal structures of S. aureus FtsZ monomers (PDB IDs: 5MN4 (T) and 5MN6 (R1) (*18*)). The R2 model was constructed directly from the E. coli FtsZ crystal structure (PDB ID: 6UNX (*38*). D) The predicted inter-monomer interface structure for each model with the interface regions labelled. Models T and R1 are shown bound to GDP and model R2 is shown bound to GTP.

However, to date, the T state has only been crystallized from *Staphylococcus aureus* FtsZ and the evidence that it is a conserved, active state is minimal. Notably, the inhibitor used to stabilize it in *S. aureus*—PC190273—does not induce polymerization in *E. coli* FtsZ (*37*) and two recently published *E. coli* crystal structures were both in the relaxed or “R” state (*38*)(*39*). The goal of this study is to determine whether the monomers within active, GTP-bound *E. coli* FtsZ filaments are in the T or R state. To do this, we use magic angle spinning (MAS) NMR under cryogenic DNP conditions (100 K) to directly observed GTP-bound, full length EcFtsZ filaments at atomic resolution. By focusing on the monomer-monomer interface within the filament, we are able to distinguish between T state monomers and R state monomers and determine that GTP-bound EcFtsZ filaments are composed primarily of T state monomers.

Magic angle spinning (MAS) NMR is an important structural technique that is well suited for the study of protein oligomers. MAS NMR has contributed greatly to our understanding of the structure of amyloid fibrils (*40*), membrane proteins (*41*), and other difficult to crystallize, insoluble protein systems (*42*). MAS NMR allows for the measurement of both through-bond and through-space signals mediated by the nuclear dipolar coupling. Experiments such as REDOR (*43*) and TEDOR (*44*) reintroduce the heteronuclear dipolar coupling allowing for the measurement of internuclear distances (*45*)(*46*). Modern multi-dimensional experiments such as z-filtered TEDOR (ZF-TEDOR) (*47*), double-REDOR (*48*), and PAIN-CP (*49*) are routinely used to measure the distance restraints used in solving protein structures (*40*)(*50*)(*51*).

The emergence of dynamic nuclear polarization (DNP) as an accessible MAS NMR technique has increased the viability of such long-range correlation experiments by drastically reducing the spectrometer time required to collect such spectra (*52*)(*53*). DNP uses cryogenic temperatures, nitroxide biradicals (or other paramagnetic centers), and high-powered microwave radiation to greatly enhance NMR signals, often on the order of 70–100x for typical proteins (*54*)(*55*)(*56*) (*57*)(*58*)(*59*) (*60*). The large increase in signal strength greatly decreases the amount of signal averaging needed to collect high quality spectra, and the low temperatures reduce protein dynamics, often leading to an increase in the number of observable resonances (*52*). Line broadening, particularly in the ^15^N dimension is a significant issue however, that has limited the usefulness of DNP in many systems (*61*).

While large proteins (>200 residues) remain difficult to study using traditional MAS NMR methods that rely on full spectral assignments, the availability of homology modeling, structural information from crystal structures of monomers, and modern spectroscopy techniques allow for targeted biological questions to be asked in the absence of full spectral assignments. Such strategies also broaden the applicability of MAS NMR to relevant and interesting biological questions by reducing the reliance on costly and difficult assignments. FtsZ is one such large protein. EcFtsZ has 383 total residues, about 300 of which are structured. Full spectral assignments for EcFtsZ monomers is not practical given the degree of spectra overlap. However, the proliferation of FtsZ crystal structures makes FtsZ a good target for exploring the applicability of MAS NMR and DNP on large, biologically interesting systems.

## Materials and Methods

### Protein Expression and Purification

*E. coli* FtsZ was expressed from a pET21b(+) plasmid generously provided by Professor Anuradha Janakiraman at the City College of New York. All protein was expressed using BL21(DE3) chemically competent cells using 0.5 mM IPTG. Isotopically enriched, protonated FtsZ was expressed using a 4:1 LB:M9 transfer with precursor concentrations as follows: for U-^15^N FtsZ: 1 g/L ^15^NH_4_Cl + 4 g/L natural abundance D-glucose; for the U-^15^N, U-^12^C FtsZ: 1 g/L ^15^NH_4_Cl + 2 g/L 99.9 % ^12^C D-glucose; for the sparsely labelled ^13^C FtsZ: 1 g/L [1,3]-^13^C-glycerol + 1 g/L natural abundance NH4Cl + 1 g/L natural abundance sodium bicarbonate (*51*). All isotopes were purchased from Cambridge Isotope Laboratory (Tewksbury, Massachusetts, USA). Unless otherwise noted, all biochemical reagents were purchased from ThermoFisherScientific (Waltham, Massachusettes, USA). GTP was purchased from SigmaAlrichMillipore (St. Louis, Missouri, USA).

For the deuterated EcFtsZ samples, BL21(DE3) chemically competent cells transformed with *pET21b(+):EcFtsZ* were adapted to ^2^H_2_O LB by first growing cells in protonated LB overnight, transferring 1 % of cells into LB media containing 70 % ^2^H_2_O and 30 % H_2_O, incubating for 24 hours, transferring 0.5 % of cells into fresh, 100 % ^2^H_2_O LB media, and growing cells for an additional 18 hours. Cells were then plated on solid agar plates containing 100 % ^2^H_2_O LB and grown for 48 hours at 37 °C. 6 isolated colonies where picked from the plate and used to establish glycerol stocks in 100 % ^2^H_2_O LB.

Partially deuterated EcFtsZ was expressed using M9 adaptation rather than a 4:1 transfer to reduce the amount of ^2^H_2_O (99.8 %) consumed. As deuteration was desired primarily at the C*α* and C*β* positions, only fractional deuteration was necessary, allowing protonated precursors to be used in a highly deuterated background media (*62*)(*63*)(*64*). Samples were prepared by growing 100 % ^2^H_2_O LB adapted BL21(DE3) cells transformed with *pET21b(+):EcFtsZ* in a 4 mL culture of ^2^H_2_O LB for 12 hours. 500 mL of 100 % ^2^H_2_O M9 media was inoculated with 1 % of the overnight cultures and grown to OD_600_ of ~0.6 in an incubating shaker at 37 °C/250 rpm, which took 18–20 hours. Expression was induced with 0.5 mM IPTG in ^2^H_2_O and expression was halted after 4 hours. For the ^2^H_2_O, 2-^13^C-glycerol EcFtsZ the M9 was prepared with 2 g/L 2-^13^C-glycerol, 1 g/L natural abundance NH_4_Cl, and 2 g/L ^13^C-sodium bicarbonate. For the ^2^H_2_O ^13^C*α*-glycine, ^13^C*β*-alanine EcFtsZ the M9 was prepared with 4 g/L ^12^C-glucose (99.9 %, Cambridge Isotopes Laboratories), 1 g/L natural abundance NH_4_Cl, 1 g/L ^13^C*α*-glycine, 1 g/L ^13^C*β*-alanine, 0.1 g/L natural abundance L-isoleucine, and 0.2 g/L natural abundance *α*-ketoisovalerate.

EcFtsZ was purified using ammonium sulfate precipitation and calcium sedimentation, as described elsewhere (*65*)(*14*).

Biochemical characterization and polymerization assays are described in the supplementary materials.

### DNP

The samples used in these experiments were U-^15^N FtsZ + [1,3]-^13^C-glycerol FtsZ polymerized together with GTP dissolved in D_2_O, U-^15^N FtsZ polymerized with GTP (^15^N control), U-^15^N, U-^12^C FtsZ + [1,3]-^13^C-glycerol FtsZ polymerized together with GTP, U-^15^N, U-^12^C FtsZ + 2-^13^C-glycerol FtsZ polymerized together with GTP, and U-^15^N, U-^12^C FtsZ + ^13^C*α*-Gly, ^13^C*β*-Ala FtsZ polymerized together with GTP. All samples were polymerized in DNP buffer (50 mM MES/NaOH pH 6.5, 50 mM KCl, 20 mM MgCl_2_, 2 mM EDTA) diluted to 30 % U-^12^C, U-^2^H glycerol bringing the final buffer concentrations to approximately 5 mg FtsZ, 30 mM KCl, 12 mM MgCl_2_, 1.2 mM EDTA, and 3 mM GTP, with 10 mM AMUPol with a final H_2_O/D_2_O ratio of approximately 65/35. Samples were centrifuged into Bruker 1.9 mm low temperature rotors and promptly frozen in liquid nitrogen to preserve stable filaments.

DNP experiments were performed on a 600 MHz (14.1 T) Bruker Avance III-DNP system at the New York Structural Biology Center (NYSBC), equipped with a 395 GHz gyrotron and an HCN triple channel probe. Additional DNP experiments were performed on a 600 MHz (14.1 T) Bruker Avance III-DNP system located at the Bruker Biospin Corporation in Billerica, Massachusetts, equipped with a 395 GHz gyrotron and an HCN triple channel probe (Bruker BioSpin, Billerica, MA). All pulse sequences used in this paper are publicly available and can be found at http://comdnmr.nysbc.org/comd-nmr-dissem/comd-nmr-solid. The CP90 pulse sequence was used for all CP spectra and the zTEDOR pulse sequence was used for all collected TEDOR spectra.

Experiments were collected with a spinning frequency, *ω_r_* /2*π*, of 14,000 ± 10 Hz and an estimated temperature of 115 ± 2.5 K. Standard *π*/2 pulse lengths of 1.48, 3.0, and 4.2 *μ*s were used for the ^1^H, ^13^C, and ^15^N channels, respectively, corresponding to *ω*_1_ /2*π* = 169 kHz (^1^H), 83 kHz (^13^C), and 60 kHz (^15^N). Heteronuclear recoupling was done using z-filtered TEDOR (ZF-TEDOR) (*47*) using mixing times between 2.0 and 24.28 ms. ^1^H decoupling of *ω*_1,*H*_/2*π* > 70 kHz was performed using SPINAL-64 decoupling (*66*).

Experiments done at Bruker BioSpin (Billerica, MA) were collected with a spinning frequency, *ω_r_*/2*π*, of 14,000 ± 10 Hz and an estimated temperature of 115 ± 2.5 K. Standard *π*/2 pulse lengths of 1.2, 3.0, and 4.15 *μ*s were used for the ^1^H, ^13^C, and ^15^N channels, respectively, corresponding to *ω*_1_/2*π* = 208 kHz (^1^H), 83 kHz (^13^C), and 60 kHz (^15^N). Heteronuclear recoupling was done using z-filtered TEDOR (ZF-TEDOR) (*47*) using mixing times between 2.0 and 24.28 ms. ^1^H decoupling of *ω*_1,*H*_/2*π* > 80 kHz was performed using SPINAL-64 decoupling (*66*).

Spectra were processed using TopSpin (Bruker BioSpin). The ^13^C chemical shifts were referenced via the substitution method to the reported chemical shifts of proline. To account for low signal-to-noise, peaks were picked by overlaying 1D ZF-TEDOR spectra and identifying peaks that occured in at least 2 spectra. Data was analyzed in python using the NMRglue library (*67*). All analysis code is available at https://github.com/kmmccoy/ZFTEDOR_fitting.

### Homology Model Construction

EcFtsZ dimer models were constructed using the Schrödinger Maestro Suite (Schrödinger Release 2020-4: Maestro, Schrödinger, LLC, New York, NY, 2020). A dimer was inferred from the *Staphylococcus aureus* crystal structure by choosing the symmetry mate that best resembled a filament for the tense (PDB ID: 5MN4) and relaxed (PDB ID: 5MN6) forms (*18*). The dimers were then templated against the wild type EcFtsZ sequence to construct homology models of the tense and relaxed (“R1”) dimers. A second relaxed dimer model (“R2”) was constructed using the *E. coli* FtsZ crystal structure (PDB ID: 6UMX) (*38*). The interface was inferred by choosing the symmetry mate that best resembled a filament. The chemical shifts for all three models were predicted using SHIFTX2 (*68*).

### Curve Fitting

In order to extract internuclear distances, the build-up curves of the identified peaks were fit using numerical simulations generated by SPINEVOLTION (*69*). The SPINEVOLUTION model included an isolated C–N pair. The t_2_ value was varied and 10 Hz of *J_cc_* coupling was included, although the fitting was found to be insensitive to the *J_cc_* values. The data was normalized to the transfer efficiency–i.e. the percentage of the CP transfer for each peak. The integrated intensity for each peak in each spectrum was divided by the integrated intensity for the same region of the CP spectrum and multiplied by 100.

The RMSE was calculated for the fits along with the *RMSE*/*σ_data_* and the 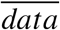—the coefficient of variation of the RMSE—as measures of goodness of fit. A good fit is defined here as a fit with a value of the *RMSE/σ_data_* of < 0.5—i.e. the standard deviation of the residuals between the fit and the data (the RMSE) is less than half of the standard deviation of the data. If this value was over 0.5 across all fitting models the curve was deemed too noisy for accurate fitting and was excluded. The fit error was estimated using jackknife error estimation.

## Results and Discussion

### Modelling The EcFtsZ Interface

Two distinct structural conformations of the FtsZ monomer have been identified in *S. aureus*: a relaxed (“R”), low energy conformation that corresponds to the majority of previously identified FtsZ monomer structures and a tense (“T”), high energy conformation that has been posited to represent the active, polymerized form of the monomer (*18*)(*31*). As compared to the R state, the T state monomer’s C-terminal subdomain is rotated and moved away from the N-terminal subdomain by nearly 30°s and is characterized by an open interdomain cleft (*18*). In the context of the filament, the T state monomer is aligned in the crystal in a manner that appears to create a straight filament with the characteristic 44 nm monomer spacing seen in EM images of straight filaments (*18*) (figure 1). However, inhibitor binding, mutagenesis, or the use of GTP mimetics must be used to trap the monomer in the T state for crystallization (*31*)(*18*)(*34*). Additionally, the T state has never been directly observed outside of *S. aureus*.

The active form of FtsZ is the filament. Thus, the active form of the monomer is the form that is present in the active (straight) form of the filament. Assuming that the two major forms of the monomer are the T and R states, we set out to identify which state is present in *E. coli* FtsZ filaments. FtsZ forms single-stranded filaments composed of a single monomer. This means that the monomer-monomer interface is unique to the filament and can, in principle, be used to characterize the active monomers. In order to understand the differences in the monomer-monomer interface between the T and R states, we constructed three separate models of the EcFtsZ dimer. We chose the dimer because it is the smallest unit of the filament containing an interface. The T state dimer model was generated by defining a dimer based on the crystal contacts in the PDB file of the SaFtsZ T state monomer (PDB ID 5MN4 (*18*)). This dimer was then templated against the EcFtsZ sequence and used to construct a homology model of the EcFtsZ dimer using Schrödinger Maestro (Schrödinger, LLC, New York, NY, 2020) (*70*). This same method was used to construct an R type dimer (model “R1”) (PDB ID 5MN6 (*18*)). Additionally, the recent EcFtsZ monomer structure (PDB ID 6UNX (*38*)) was used to define a second R state dimer (model “R2”) based on the crystal contacts present in the PDB file (figure 1).

While the interfaces in the three models are similar, the T model interface is substantially larger than the other two in that it contains more interface residues. We defined an interface residue as any residue with at least one carbon atom within 5 Å of the other monomer. Using this definition, the T model has 78 interface residue, the R1 model has 41 interface residues, and the R2 model has 38 interface residues. The majority of the R1 and R2 interface residues are in what we are terming the hinge region of the interface (figure 1). The hinge region consists of the tip of H7 (on the plus end of the monomer) and the flexible loop between H7 and H6 on the plus end and H10 on the minus end (see (*38*) and figure 1D for numbering scheme). The hinge region is enriched in hydrophobic residues (particularly leucine and valines) and while the exact contacts differ between interface models, the hinge contains close contacts between the monomers in all three models. The T model’s additional interface residues are present across all interface regions, including the T7 loop and the GTP-binding pocket. Additionally, many interface residues in the T model are located in the clasp region. The clasp consists primarily of the S3 and H3 on the plus end of the monomer. In the T model, this region makes contacts with the T7 loop and H8 in the C-terminal subdomain. In the R1 model the clasp region does not make contact with the minus end of the neighboring monomer. In the EcFtsZ crystal structure reported by Schumacher et al. (*2020*) (*38*) (the R2 model) this loop extends across the interface and does make contact with part of the T7 loop, however the contacts are substantially different than those in the T model (figure 1D).

### ^13^C Natural Abundance Background in Large Proteins

We used this information to design a set of solid-state MAS NMR experiments to probe the interface in active, GTP bound EcFtsZ filaments that allowed us to determine which interface is present in the the filaments. MAS NMR is well suited for this question because it allows us to spectroscopically isolate the inter-monomer interface within the filaments using well established dipolar recoupling techniques. It also allows us to study active, GTP bound filaments composed of full-length, wild-type protein. Unlike X-ray crystallography, where filament structure must be inferred from crystal packing, MAS NMR allows us to study the filaments directly at atomic resolution. However, MAS NMR is limited by low sensitivity and high noise. Long-range correlation experiments of the type used here are particularly low signal-to-noise, with as little as 2% transfer efficiency common for 4–5 Å contacts (*52*) (figure 2A). We used dynamic nuclear polarization (DNP) to enhance our signal intensity an average of 70-fold. We were able to exploit DNP’s need for cryogenic sample temperatures (often been 100–120 K) to study full-length, WT EcFtsZ bound to GTP rather than a slowly hydrolyzing GTP-mimetic such as GMPCPP. By freezing our samples in liquid nitrogen immediately after the addition of GTP (there was on average 5–10 minutes between the addition of GTP to our samples and the immersion of a packed sample container into liquid nitrogen) we trapped GTP-bound filaments in their active state for the duration of our experiments.

**Figure 2:**
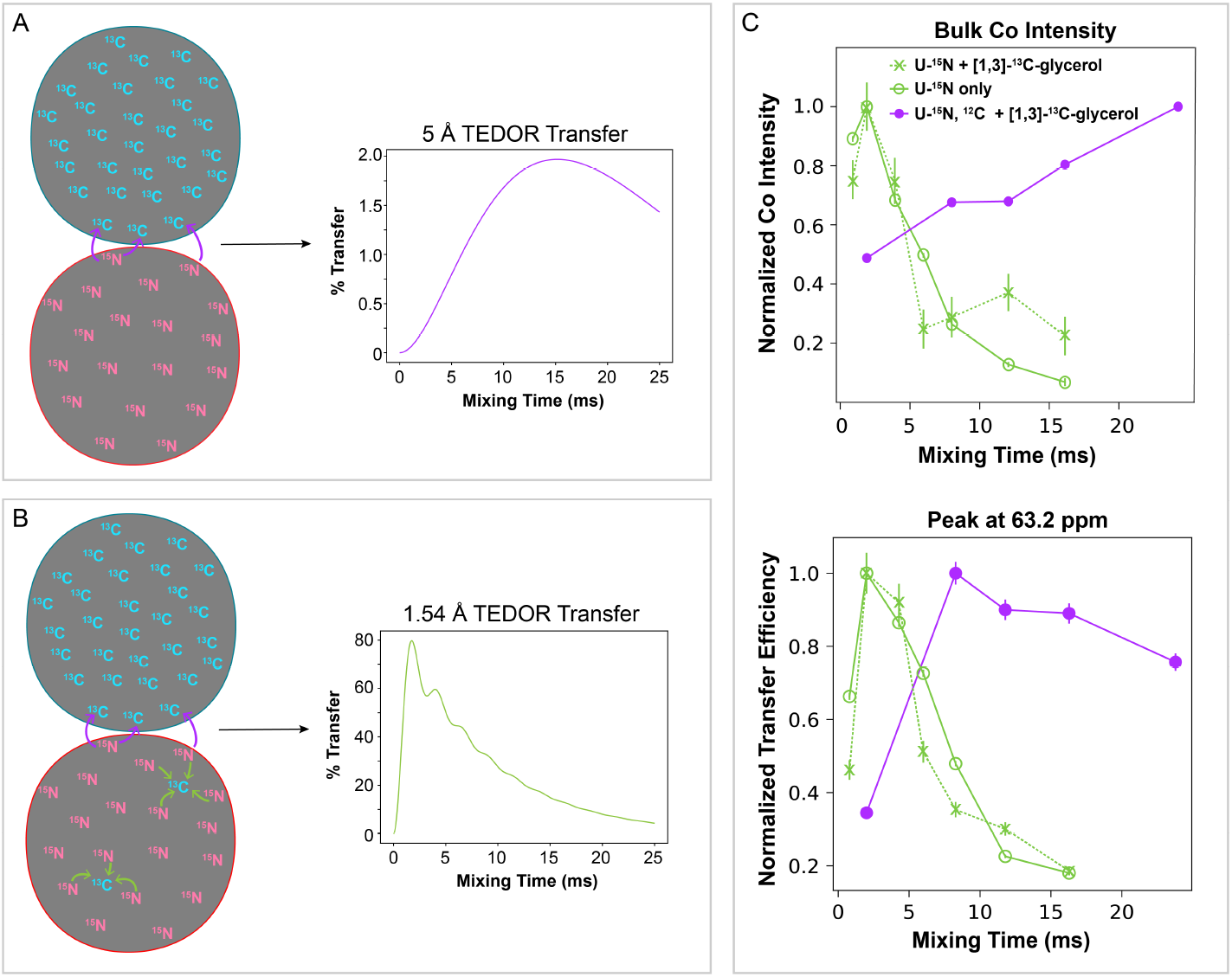
A) Schematic of the differential isotopic labelling scheme used in this study. FtsZ monomers isotopically enriched only with ^15^N were mixed with monomers enriched only with ^13^C and polymerized together, forming a filament with randomly incorporated monomers. A spectroscopic technique (ZF-TEDOR) was employed to isolate only the ^13^C signals from atoms that are directly adjacent to ^15^N atoms. In the case of the differentially labelled filament, this should only occur at interfaces, leading to the simulated TEDOR build-up curve shown. B) Schematic of differential isotopically labelled dimer in the presence of natural abundance ^13^C in the ^15^N labelled monomers. This changes the expected build-up curve, as shown in the simulated TEDOR curve. Both simulated TEDOR curves were calculated using an analytical expression that approximates TEDOR build-up (*71*). Parameters for the simulations were as follows: a single carbon either 1.54 Å (b) or 5 Å (a) from a single nitrogen (passive dipolar coupling of 0 Hz) with a T_2_ value of 8 ms, a scaler value of 1, and 10 Hz of second order J-coupling. C) Experimental build-up curves from differentially labelled FtsZ monomers bound to GTP for both the bulk carbonyl intensity and an individual peak. FtsZ labelled with [1, 3]-^13^C-glycerol was mixed with U-^15^N enriched FtsZ made with either natural abundance glucose (blue) or ^12^C-glucose (99.9 %) (green). FtsZ filaments composed of only U-^15^N enriched FtsZ made with either natural abundance glucose (orange) were used as a control. The curves from the filaments made with natural abundance glucose show a very similar build-up pattern to those from the U-^15^N only filaments, with build-up peaking around 2.0 ms mixing, consistent with 1 or 2 bond distances. When filaments were made using ^12^C-glucose, the polarization transfer is consistent with long distance, inter-monomer contacts.

In order to isolate the longitudinal monomer interface of EcFtsZ filaments, we prepared a DNP sample using a differential isotopic labelling approach previously employed in the structural analysis of amyloid fibrils (*72*)(*52*) (figure 2). U-^15^N FtsZ was mixed with [1, 3]-^13^C-glycerol FtsZ in 1:1 amounts, polymerized together using GTP, packed into a 1.9 mm DNP rotor, and frozen in liquid nitrogen to ensure that the majority of filaments were in their active state. ZF-TEDOR 1D experiments were run using mixing times from 0.8 ms to 16 ms. Despite the expected lack of directly bonded ^13^C–^15^N pairs due to the differential labelling scheme, the ZF-TEDOR buildup curves had signal maxima at short mixing times, corresponding to distances only achievable through a short transfer—~2–3 Å, corresponding to 2–6 ms mixing times (*47*) (figure 2).

Due to the differential labelling scheme, this is most likely from natural abundance ^13^C present in the U-^15^N monomers. In order to test this, we prepared a sample composed of only U-^15^N FtsZ, filamented in the same manner as the mixed label sample. This ^15^N control sample showed nearly identical build-up curves as the mixed label sample (figure 2), particularly at short TEDOR mixing times, demonstrating that the ZF-TEDOR spectra were dominated by intra-monomer N–C transfers between the labelled ^15^N and natural abundance ^13^C within the ^15^N labelled monomers. Interestingly, the build-up curve of carbonyl intensity in the mixed label sample differs from that of the ^15^N control sample at long mixing times (notably 12 ms and 16 ms), showing additional magnetization build-up at > 10 ms (figure 2C), indicating that inter-monomer transfer is occurring, but is obscured by the natural abundance ^13^C background.

In order to understand the magnitude of the effect of natural abundance ^13^C in the U-^15^N enriched monomers on the ZF-TEDOR spectra, it is helpful to compare the number of expected intra-monomer ^13^C–^15^N contacts to the number of ^13^C–^15^N contacts expected across the interface. In the SaFtsZ T state monomer crystal structure (PDB ID 5MN4)(*18*), there are 80–100 intermonomer ^13^C–^15^N contacts within 5 Å. Given the ~ 50 % sparse carbon labelling from [1, 3]-^13^C-glycerol (*51*), that comes out to a predicted 40–50 inter-monomer contacts across the interface. In the EcFtsZ monomer, the average backbone nitrogen is within 5 Å of 15.4 carbons. Sidechain nitrogens are, on average, within 5 Å of 8-9 carbons (with the exception of histidine’s N*δ*1, which is within 5 Å of 13.5 carbons). If we just consider the backbone, based on the ~300 structured residues in EcFtsZ, we can assume that there are 300 backbone nitrogens, each with 15.4 N–C contacts, giving a total of ~ 4,600 contacts per monomer. Given the natural abundance ^13^C of 1.1 %, that means that there are ~50 ^13^C–^15^N contacts within 5 Å per U-^15^N labelled EcFtsZ monomer. Assuming 40–50 inter-monomer contacts, this means that there are at least as many intra-monomer N–C contacts as inter-monomer ones. Given this, it is not surprising that the ZF-TEDOR spectra are dominated by natural abundance background signal.

In order to reduce the contribution from natural abundance ^13^C, we prepared a sample where the U-^15^N FtsZ was grown in media containing ^12^C-glucose (99.9 %) as the sole carbon source. The use of ^13^C-glucose significantly reduced the amount of TEDOR signal at short mixing times (figure 2C), suggesting a large reduction in the amount of intra-monomer TEDOR transfer. The ^12^C-glucose is 0.1 % ^13^C versus the 1.1 % ^13^C present in natural abundance glucose. This reduces the amount of ^13^C nuclei present in each U-^15^N monomer to ~ 1.6, allowing for the observation of unambiguous inter-monomer TEDOR transfer as measured by magnetization build-up at long TEDOR mixing times (> 16 ms) (figure 2C). This strategy is generally applicable to MAS NMR studies of large proteins with small interface surfaces. It is likely that when combined with spectroscopy techniques such as an additional short-mixing time REDOR block (*73*)(*74*)(*75*), this will allow the removal of the majority of background signal from natural abundance ^13^C.

### Characterizing the EcFtsZ Interface

Three samples of EcFtsZ filaments were prepared using different isotopic labelling schemes. [1, 3]-^13^C-glycerol + U-^15^N, ^12^C mixed label EcFtsZ (referred to subsequently as the 1, 3-glycerol sample) and 2-^13^C-glycerol + U-^15^N, ^12^C mixed label EcFtsZ (referred to as the 2-glycerol sample) were used to get broad coverage of interface residue atoms. Additionally, we prepared ^13^C*α*-glycine and ^13^C*β*-alanine labelled EcFtsZ mixed with U-^15^N, ^12^C EcFtsZ to reduce spectral crowding (referred to as the ^13^C*α*-Gly, ^13^C*β*-Ala sample). Additionally, the 2-glycerol and ^13^C*α*-Gly, ^13^C*β*-Ala samples were fractionally deuterated on the ^13^C enriched monomers in order to obtain suitable T_2_ values for the collection of long mixing time spectra. For each sample we collected a series of 1D ZF-TEDOR spectra at various mixing times and observed polarization build-up in a distance dependant manner. See figure S4 for representative spectra from each sample.

Peaks were picked for each sample by identifying peaks that appeared in at least two of the ZF-TEDOR spectra. Due to extremely low signal-to-noise (S/N), it was difficult to identify non-noise peaks in each spectrum. However, it is unlikely that a noise peak will occur at the same location in two separate spectra, thus we reasoned that peaks that appear in the same location in multiple spectra are likely to not be due to noise. Peak locations were identified using the peak maxima; the spectra were too S/N limited to accurately apply line shape fitting. This method led to the identification of 49 peaks in the 1, 3-glycerol spectra, 83 peaks in the 2-glycerol spectra, and 94 peaks in the ^13^C*α*-Gly, ^13^C*β*-Ala spectra. ZF-TEDOR build-up curves were plotted for each peak by integrating a slice of the spectrum 1 ppm wide centered at the peak. Transfer efficiencies were calculated by dividing the integrated intensity of each peak by the integrated intensity of the same slice of the ^1^H–^13^C cross polarization spectrum. This procedure was repeated for the same regions of the U-^15^N sample spectra as a control. Figure 1C shows a representative build-up curve (see Figure S3 and S4 for more build-up curves). The transfer efficiencies have maxima a longer mixing times (> 12 ms), which is consistent with internuclear distances between 4–6 Å, indicating that the contacts being observed are inter-monomer.

Due to the lack of full resonance assignments for EcFtsZ, along with low S/N in the ZF-TEDOR spectra that only allow for the collection of 1D spectra, full characterization of the EcFtsZ interface is not possible with the data set described here. However, using the dimer models described above, we were able to generate predicted chemical shift lists to compare to the peaks present in the spectra. We predicted the chemical shifts of all three dimer models using SHIFTX2 (*68*) and extracted the chemical shifts of residues present as the interface. In order to test the experimental data against the interface models, we identified predicted chemical shifts within ±0.5 ppm of each experimental peak and compared the predictions. A cut of ±0.5 ppm was used because it corresponds to the average accuracy of ^13^C chemical shift prediction using SHIFTX2 (*68*). Additionally, it corresponds to our estimated ^13^C line width.

We ranked each model’s prediction based on whether the predicted residue appeared at the interface, if it was expected to be labelled, or if it appeared on a lateral surface and may be explained by lateral contacts between bundled filaments. Most peaks were equally well explained by all three models—meaning that all three models had predicted chemical shifts within 0.5 ppm of the experimental peak that correspond to labelled atoms in interface residues. Predicted shifts from residues appearing at the interface and atoms that are expected to be labelled at > 50 % efficiency were ranked the highest, followed by labelled atoms appearing at a lateral surface, unlabelled atoms that do not appear at an interface, and finally, peaks that cannot be explained by any predicted chemical shift.

Most peaks can be explained equally well by any of the three models. However, a more telling criterion is how many peaks each model fails to adequately account for. Table 1 summarizes this analysis. Of the 219 total peaks identified across all three samples, the T model can account for all but 24 of them. The R1 accounts for all but 58 peaks, and the R2 model fails to account for 75 peaks. 5 additional peaks in the 2-glycerol spectra cannot be explained by any model used in the analysis. With the exception of a peak at 170.4 ppm, these peaks (135.7 ppm, 140.9 ppm, 156.7 ppm, and 158.3 ppm) correspond to low S/N regions and may be due to noise. Alternatively, they could result from side-chain conformations not seen in the crystal structures used to construct the models.

**Table 1:** Model Comparison By Sample

| Sample | Experimental Peaks | Not Explained By T Model | Not Explained By R1 Model | Not Explained By R2 Model |
| --- | --- | --- | --- | --- |
| 1,3-glycerol | 50 | 3 | 14 | 17 |
| 2-glycerol | 75 | 10 | 16 | 22 |
| $^{13}\text{C}\beta\text{-Ala}$ $^{13}\text{C}\alpha\text{-Gly}$ | 94 | 11 | 28 | 36 |

We also identified 29 peaks that are accounted for by only the T model or only an R model (table 2). In some cases, the experimental peak is within 0.5 ppm of two predicted chemical shifts, in which case the peak may correspond to overlapping peaks. We included peaks that may correspond to either a unique peak or two peaks in a single model, so long as they were not well explained by the other models.

**Table 2:** Model Comparison By Peak

| Peak (ppm) | Predicted Assignment | Model | Sample | Tolerance (ppm) |
| --- | --- | --- | --- | --- |
| 14.9 | Ile200 Cδ1 | T | 2-glycerol | 0.6 |
| 20.3 | Thr201 Cγ2 | R1 | 2-glycerol | 0.9 |
| 20.3 | Val171 Cγ2 | R1 | $^{13}\text{C}\beta$-Ala, $^{13}\text{C}\alpha$-Gly | 0.65 |
| 22.1 | Thr201/Thr296 Cγ2 | T | 2-glycerol | 0.6 |
| 22.4 | Ala284/Ala286 Cβ | R1 | $^{13}\text{C}\beta$-Ala, $^{13}\text{C}\alpha$-Gly | 0.65 |
| 26.0 | Leu169 Cδ1 | T | $^{13}\text{C}\beta$-Ala, $^{13}\text{C}\alpha$-Gly | 0.7 |
| 26.4 | Ile294 Cγ1 | T | 1,3-glycerol | > 1.0 |
| 32.9 | Val171/Val262 Cβ | R1 | 2-glycerol | 1.0 |
| 33.4 | Lys51/Met206 Cβ | T | 1,3-glycerol | > 1.0 |
| 43.1 | Leu168 Cβ | T | 2-glycerol | 0.6 |
| 48.2 | Gly20 Cα | T | $^{13}\text{C}\beta$-Ala, $^{13}\text{C}\alpha$-Gly | 0.6 |
| 53.7 | Arg202 Cα | T | 1,3-glycerol | 0.75 |
| 57.2 | Ser287/Ser177 Cα | R1/R2 | $^{13}\text{C}\beta$-Ala, $^{13}\text{C}\alpha$-Gly | > 1.0 |
| 57.6 | Ser287/Ser177 Cα | R1/R2 | $^{13}\text{C}\beta$-Ala, $^{13}\text{C}\alpha$-Gly | > 1.0 |
| 59.9 | Thr296 Cα | T | $^{13}\text{C}\beta$-Ala, $^{13}\text{C}\alpha$-Gly | 1.0 |
| 61.8 | Ser177/Thr65 Cα | R1/R2 | $^{13}\text{C}\beta$-Ala, $^{13}\text{C}\alpha$-Gly | 0.55 |
| 63.2 | Pro134 Cα/Ser177 Cβ | R1 | 1,3-glycerol | 0.55 |
| 63.3 | Thr65 Cα | T | $^{13}\text{C}\beta$-Ala, $^{13}\text{C}\alpha$-Gly | > 1.0 |
| 63.5 | Ile200 Cα | R1/R2 | 2-glycerol | 0.65 |
| 64.6 | Thr201 Cα | T | $^{13}\text{C}\beta$-Ala, $^{13}\text{C}\alpha$-Gly | > 1.0 |
| 66.4 | Thr215 Cα | T | $^{13}\text{C}\beta$-Ala, $^{13}\text{C}\alpha$-Gly | > 1.0 |
| 67.5 | Thr281 Cα | T | $^{13}\text{C}\beta$-Ala, $^{13}\text{C}\alpha$-Gly | 0.6 |
| 72.5 | Thr296 Cβ | T | 1,3-glycerol | > 1.0 |
| 158.8 | Arg202 Cz | T | 2-glycerol | 0.8 |
| 160.8 | Arg174/Arg271 Cz | R1 | 2-glycerol | 0.6 |
| 181.5 | Glu250 Cδ | T | 1,3-glycerol | > 1.0 |

Of the 29 peaks identified, 18 (62 %) are best explained only by the T model, 6 (21 %) are best explained only by the R1 model, and none are best explained by the R2 model. However, 5 (17 %) are best explained by either the R1 or the R2 model, but not the T model. Several of the predicted assignments correspond to atoms in the same residue. For example, the peak at 53.7 ppm in the 1, 3-glycerol spectra can be explained by Arg202’s C*α* and the peak at 158.8 ppm in the 2-glycerol spectra can be explained by Arg202’s Cz. Similarly, Thr296 can explain the peaks at 59.9 ppm in the ^13^C*β*-Ala, ^13^C*α*-Gly spectra (C*α*), 72.5 ppm in the 1, 3-glycerol spectra (C*β*), and potentially the peak at 22.1 ppm in the 2-glycerol spectra (C*γ*2). The recurrence of the same residues across all 3 samples demonstrates that the same interface form is present across samples.

Table 2 also lists the tolerance of the model assignments—defined here as the maximum ppm window for a single model assignment. For example, for the peak at 14.9 ppm, if a ± 0.5 ppm window is used in the analysis, only the T model explains the peak well. However, if a ± 0.6 ppm window is used, both the T and the R2 models can explain the peak. The tolerances show that the result of this analysis does not substantially change if we use larger windows. Using a window of ± 0.6 ppm, there are 26 peaks, 17 of which are best explained by the T model (65 %), 5 of which are explained only the R1 model (19 %), and 4 of which are best explained by either the R1 or the R2 model but not the T model (15 %). With a ± 0.8 ppm window, 8 of the resulting 13 peaks are best explained by the T model (62 %), 2 are best explained by the R1 model (5 %), and 3 are best explained by either the R1 or the R2 model (23 %).

The fact that the T model better accounts for the experimental data, regardless of the size of the ppm window used, is evidence that the EcFtsZ filaments are primarily composed of T monomers. That some peaks are not well explained by the T model used in this analysis suggests that either some percentage of interfaces present in the sample contain R state monomers—either due to filaments with mixed state or to non-polymerized oligomers—or that the physiological interface differ from the models used for this analysis. With continued refinement of the model, and subsequent rounds of chemical shift prediction and analysis, we expect that more peaks will be well explained by the model.

To build on this analysis, we took the residues that correspond to unique assignments and identified which region of the interface they are located in (table 3). Glu250 and Thr281 do not occur at the interface. Rather, they are both located on the lateral surface of the monomer and likely represents lateral contacts between bundled filaments. Gly20 is located in the H1 helix and is part of the GTP-binding pocket. Leu168, Leu169, and Val171 are all in the loop region between H6 and H7 and form the core of the hinge region on the plus end of the monomer. Ile294 and Thr296 are also part of the hinge region on the minus end. They are both located in the S9 strand, which, due to the orientation of the C-terminal subdomain in the T conformation, faces H6 and the helical loop between S5 and H5 on the plus end of the interface. Close inspection of the models shows that in the R state models, S9 is rotated so that is faces away from the hinge, and thus, contacts are not expected from this region (figure 1). Figure 3 illustrates the location of predicted and observed contacts in the T model and shows the difference in the T and R1 interface contacts.

**Table 3:** Interface Residue Locations

| Residue | Model | Region |
| --- | --- | --- |
| Gly20 | T | GTP-binding pocket |
| Thr65 | T | Clasp |
| Leu168 | T | Hinge |
| Leu169 | T | Hinge |
| Val171 | R1 | Hinge |
| Ile200 | R1/T | T7 loop |
| Thr201 | R1/T | T7 loop |
| Arg202 | T | T7 loop |
| Thr215 | T | Clasp |
| Glu250 | T | Lateral Contact |
| Thr281* | T | Lateral Contact |
| Ile294 | T | Hinge |
| Thr296 | T | Hinge |

\*The peak at 67.5 ppm may also correspond to Thr277 C $\alpha$ , which occurs in the Hinge region of the interface

**Figure 3:**
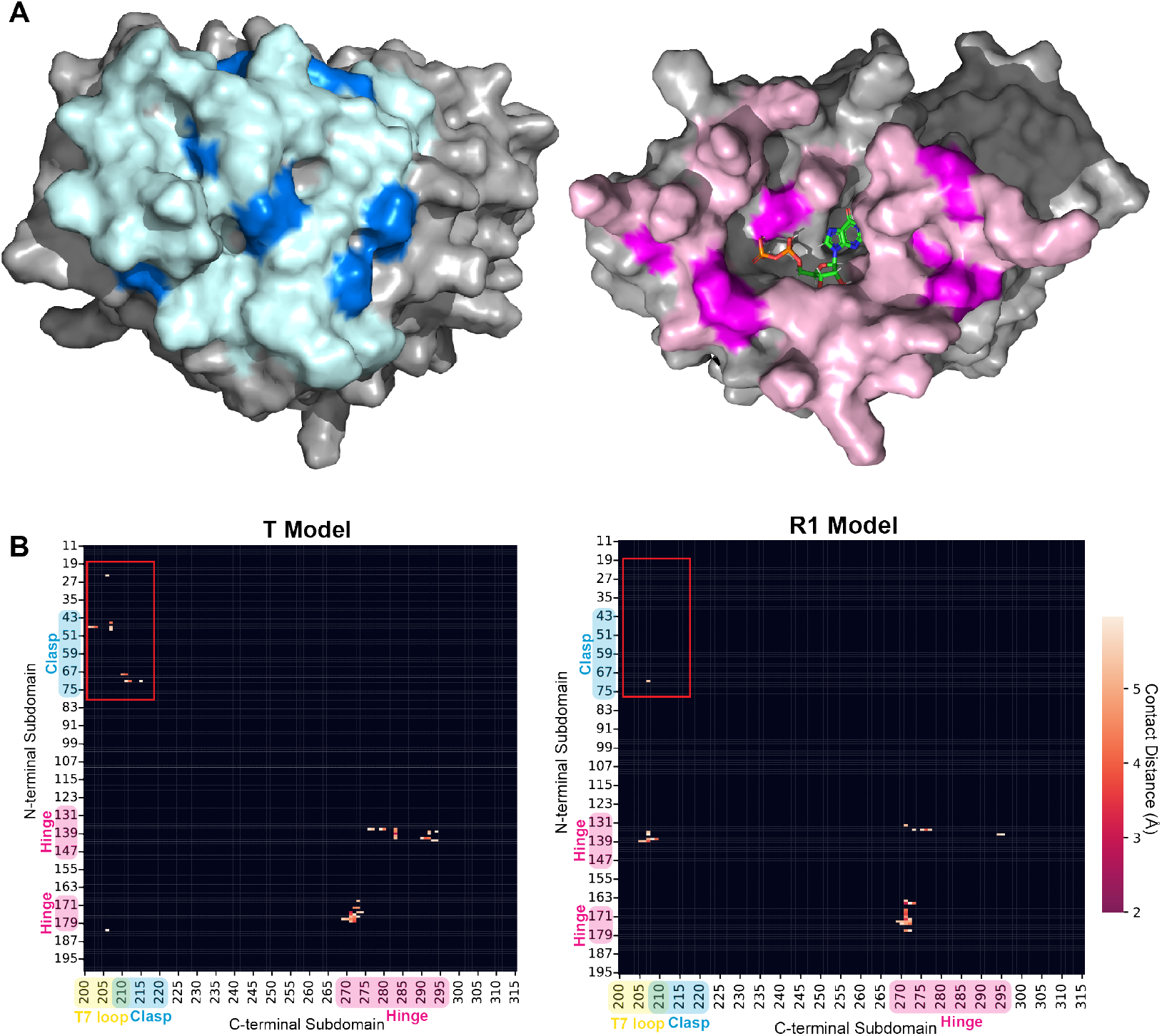
A) EcFtsZ tense model dimer structure generated using homology modelling in Schrödinger Maestro templated against PDB ID 5MN4 (*18*). The location of the predicted interresidue contacts are shown in light pink and light blue, and the location of observed contacts are shown in dark pink and dark blue. B) Contact map of the predicted T and R1 dimer interfaces showing sub-6 Å contacts between heavy atoms. Clasp region contacts are highlighted (red box) showing the presence of such contacts in the T model and absence in the R1 model.

The clasp region and its contacts with the T7 loop are also present in the data. Ile200, Thr201, and Arg202 are in the T7 loop at the very end of H7. Ile200 and Thr201 are also expected to have contacts in the R models, but Arg202 is not because the T7 loop is shifted away from the clasp in the R conformations. Thr65 is located in the clasp region on the plus end of the interface and is predicted to have an interface contact in the R2 model as well as the T model. However, the difference in the loop conformations between the T and R2 models means that the chemical shift is significantly different between the two sets of predictions. In the R2 model, T65 C*α* is predicted to have a chemical shift of 61.9 ppm, whereas in the T model it is predicted to have a shift of 66.1 ppm, which aligns with the observed shift of 66.4 ppm. Thr215 is present in the minus end of the interface in the H8 helix (figure 1; figure 4). In the R models, H8 is shifted up and away from the clasp region of the plus end interface and Thr215 does not make have contacts, and its presence in the data is a good indicator that the T state interface is being observed.

**Figure 4:**
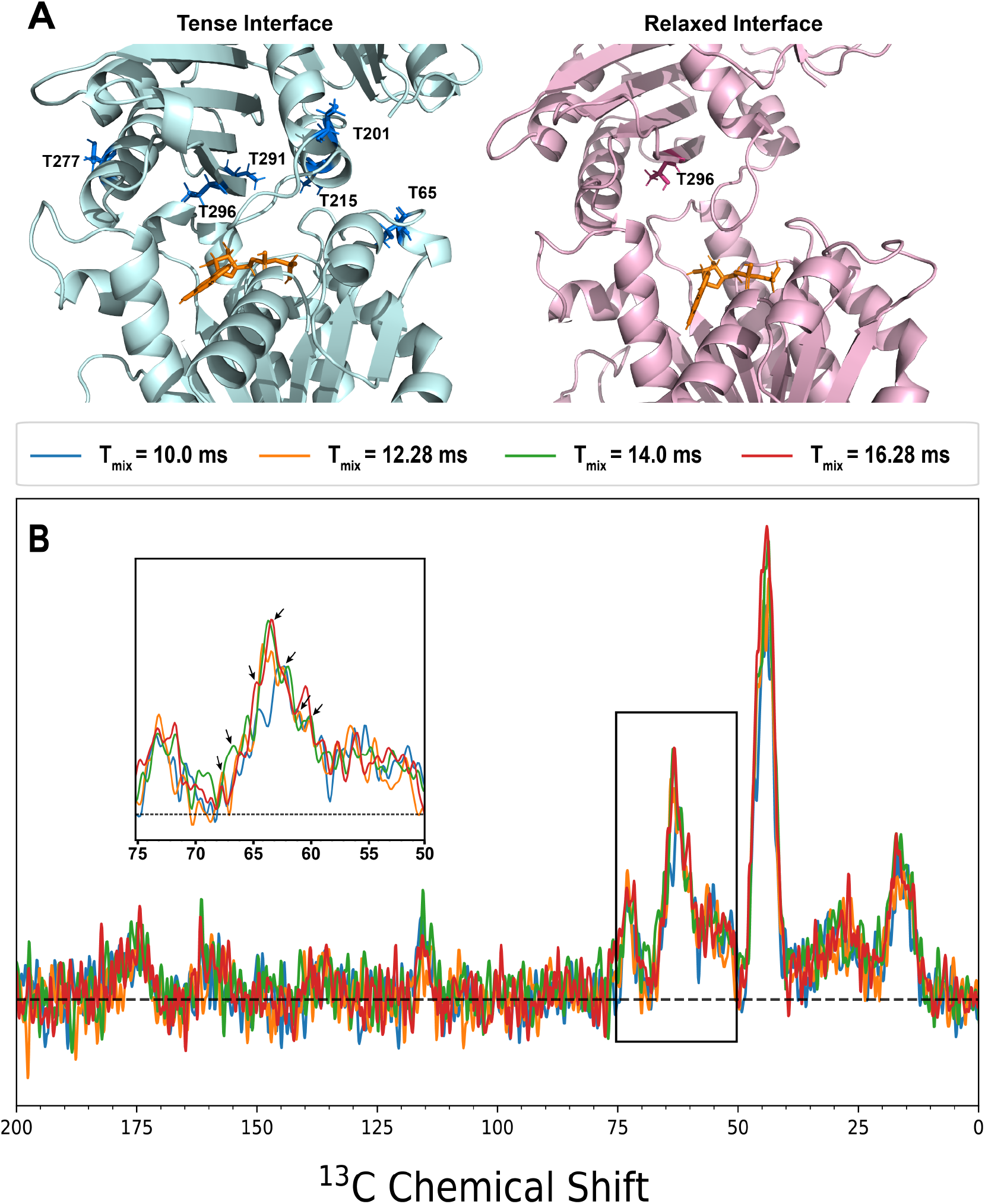
A) Location of threonine residues at the interface in the T and R1 interface models. In the T model, six threonine residues are located within 5 Å of the interface. However, in the R1 model, structural rearrangement of the C-terminal subdomain moves all but one threonine away from the interface (Thr296). B) ^13^C*β*-Ala, ^13^C*α*-Gly ZF-TEDOR spectra for mixing times of 10.0 ms (blue), 12.28 ms (orange), 14.0 ms (green), and 16.28 ms (red) showing significant build-up between 60–70 ppm. All spectra collected with 5120 scans and processed with 50 Hz of exponential linebroadening. Inset: a close up of the region between 60–75 ppm with arrows indicating the 7 identified unique peaks.

### Interface Threonine Shifts Indicate a T Monomer State

In general, threonines are good indicators of interface conformation. The T model interface contains 6 unique threonines (Thr65, Thr201, Thr215, Thr277, Thr291, and Thr296), whereas the R1 interface contains only 1 (Thr 296), and the R2 interface only contains Thr65 (figure 4A). In addition, threonine chemicals shifts are easily distinguished from other residue types, making it a good marker for EcFtsZ interface conformation. While threonines can be difficult to specifically isotopically enrich due to isotope scrambling from the bacterial metabolism, we can exploit those same amino acid metabolic pathways and scrambling. For example, glycine C*α* scrambles to serine and from serine to threonine (see supplemental materials for more detail.) Because of this, threonine C*α*s were apparent in our ^13^C*β*-Ala, ^13^C*α*-Gly sample spectra (Figure S1). While there is some overlap in the chemical shifts of serine C*α*s with threonine C*α*s, there are only, at max, 2 serines expected to be at the interface in any of the models (1 in the T model, 2 in the R1 model, and 1 in the R2 model), so serine C*α* is not expected to contribute significantly to peaks arising in the region between ~60–65 ppm.

Figure 4B shows the ZF-TEDOR spectra for long mixing times for the ^13^C*β*-Ala, ^13^C*α*-Gly sample. There is a large set of overlapping peaks evident between 60–70 ppm. Nine peaks were identified in this region, seven of which can be attributed to individual labelled atoms based on the chemical shift predictions for the T model (table 4). Six of the seven peaks can be assigned to threonine C*α*s, with the seventh peak being the sole serine C*α* expected in the T interface. One of the peaks is assigned to Thr281, which is not expected to be present at the interface and is likely a lateral contact between neighboring filaments.

**Table 4:** Threonine Peaks in ^13^C*β*-Ala, ^13^C*α*-Gly ZF-TEDOR Spectra

| Peak (ppm) | Predicted Assignment | Peak (ppm) | Predicted Assignment |
| --- | --- | --- | --- |
| 59.9 | Thr296 $\text{C}\alpha$ | 68.9 | Thr215/Thr277/Thr291 $\text{C}\beta$ |
| 60.9 | Ser177 $\text{C}\alpha$ | 69.7 | Thr201 $\text{C}\beta$ |
| 62.3 | Thr291 $\text{C}\alpha$ | 70.6 | Thr65 $\text{C}\beta$ |
| 63.3 | Thr65 $\text{C}\alpha$ | 71.7 | Thr162/Thr291 $\text{C}\beta$ |
| 64.6 | Thr201 $\text{C}\alpha$ | 72.5 | Thr296 $\text{C}\beta$ |
| 66.4 | Thr215 $\text{C}\alpha$ | 73.2 | glycerol |
| 67.5 | Thr281 $\text{C}\alpha$ | 74.0 | glycerol |

Five of the six predicted threonine peaks can be seen in the spectra, missing only Thr277. Thr277 is predicted to have a chemical shift of 66.96 ppm, which is 0.04 ppm away from the 0.5 ppm cut-off used in this analysis, so it is possible that the peak at 67.5 ppm is from Thr277 rather than Thr281. Regardless, the presence of five of the six predicted threonine C*α* peaks in the ZF-TEDOR spectra indicates that the EcFtsZ filaments adopt a T monomer conformation similar to the T form of the SaFtsZ monomer.

Additionally, there is some signal in the region from 70–75 ppm in the ^13^C*β*-Ala, ^13^C*α*-Gly (figure 4B), which is centered around 73 ppm. While it is possible that this is from the glycerol peak expected to be at 74.9 ppm, it could also be due to an increased level of ^13^C at the threonine C*β* position over the 0.1 % ^13^C background. When threonine aldolase converts glycine to threonine the threonine C*β* comes from acetaldehyde, which is produced using pyruvate. Pyruvate, in turn, can interconvert with L-alanine. The ^13^C*β* L-alanine used to produce the same had natural abundance levels of ^13^C at all other positions (1.1 %), meaning that some pyruvate, and thus some percentage of threonine C*β*s, have a higher occurrence of ^13^C than the background 0.1 %.

In order to test this, we identified 7 peaks between 68–74 ppm (68.9 ppm, 69.7 ppm, 70.6 ppm, 71.7 ppm, 72.5 ppm, 73.2 ppm, and 74.0 ppm). The peaks at 70.6 ppm and 72.5 ppm both appear in other datasets—a peak at 70.6 ppm appears in the 2-glycerol spectra and a peak at 72.5 ppm appears in the 1, 3-glycerol spectra; additionally, a peak at 72.7 ppm appears in the 2-glycerol spectra—and can be assigned to Thr65 C*β* and Thr296 C*β* respectively. The peak at 69.7 ppm can be assigned to Thr201 C*β* and the peak at 68.9 ppm is within ± 0.5 ppm of the predicted chemical shifts of Thr215, Thr277, and Thr291 C*β*. 71.7 ppm is not within 0.5 ppm of any predicted interface threonine shifts. Interestingly, a peak at this location also occurs in the 1,3-glycerol spectra, suggesting that it is a real peak. It could be due to a lateral surface contact from Thr162 C*β*. Alternately, one of the C*β* chemical shifts from Thr215, Thr277, or Thr291 could be significantly shifted relative to the predictions. Such a chemical shift perturbation would suggest a significant structural change from the model and is likely to represent a threonine that is in a sheet rather than a helix or coil (*76*). Thr215 and Thr277 are both located in helices and are unlikely to have such a perturbation. However, Thr291 is located at the very end of sheet S9 (Figure 4A), and so it is plausible that the peak at 71.7 ppm corresponds to Thr291 C*β*.

The peaks at 73.2 ppm and 74.0 ppm cannot be explained by any predicted chemical shifts and are generally too downfield to correspond to threonine C*βs* (*76*). These peaks could very well be due to contacts between solvent glycerol and solvent exposed ^15^N atoms. It is possible that one or more of the identified peaks in the threonine C*α* region also correspond to glycerol, however it is unlikely that all or even most of them do. Taken together with the high degree of agreement between peaks in the threonine C*α* and C*β* regions and the predicted chemical shifts, it is likely that the majority of the identified peaks correspond to the assigned threonine nuclei, however more work is needed to confirm this.

### ZF-TEDOR Build-up Curve Fitting

In order to extract the dipolar coupling strength—and thus the internuclear distance—from the ZF-TEDOR build-up curves, we implemented numerical simulations using SPINEVOLUTION (*69*). We defined a spin system with a single C–N pair, due to the sparse ^13^C labelling patterns and the unlikelihood of more than one backbone nitrogen being within 5 Å of a cross-interface carbon. Experimental curves were fit using the lmfit library (*77*) with error weighted Nelder-Mead minimization (*78*). All reported error was generated using jackknife error estimation. A detailed discussion of the fitting implemented here can be found elsewhere (*79*).

For the purpose of this analysis, we selected peaks from our previous analysis that correspond to a single predicted chemical shift from the T model. We did not include peaks from the 1, 3-glycerol spectra however, due to that dataset containing fewer time points. Using this method, we identified 7 peaks in the 2-glycerol 1D ZF-TEDOR spectra and 12 peaks in the ^13^C*β*-Ala, ^13^C*α*-Gly sample spectra (table S2). Notably, four of the peaks identified in the ^13^C*β*-Ala, ^13^C*α*-Gly sample spectra are predicted to be from atoms in residues that also appear in the 2-glycerol spectra. Ile200 C*δ*1 and C*α* both appear in the 2-glycerol spectra (table 2) and Ile200 C*γ*2 appears in the ^13^C*β*-Ala, ^13^C*α*-Gly spectra. Ala72 C*α* appears in the 2-glycerol spectra and Ala72 C*β* is in the ^13^C*β*-Ala, ^13^C*α*-Gly spectra. Thr65 and Thr296 also have atoms that appear in both datasets. This demonstrates a high degree of reproducibility across samples and indicates that the same interface is present across samples.

This system gave good fits for all measured build-up curves (table 5; figure 5). We compared our measured distances to the predicted distances between each measured carbon and the nearest cross-interface nitrogen (table 5). In most cases, the T model distance was closer to our measured distance than either R model. However, even when the T model agreed best with our measurement, the model distances are generally longer than our measured distances, suggesting that the interface is tighter than it appears in crystal structures. Additionally, in the cases of Ile176 C*γ*2 and Ala72 C*α* and C*β*, the measured distances agreed better with the R1 model than the T model predictions. In the case of Ile176 C*γ*2 (17.1 ppm), the T1 model predicts a contact between Ile176 C*γ*2 and a backbone nitrogen of Arg271 with a distance of 5.4 Å. In the R1 model, Arg271 is rotated so that it’s sidechain nitrogens are facing Ile176, making a much shorter, 3.5 Å contact, which is in agreement with the measured distance of 3.5 ± 0.6 Å. In the case of Ala72, in the T model Ala72 makes contact with the backbone nitrogen of Thr215 of the opposing monomer, at about a 7 Å distance. However, in both R models, Ala72 makes contacts with the backbone nitrogens in the T7 loop (209/210/211) with distances between 5.4 and 7.8 Å. However, the agreement of the measured distances for T7 loop residues Ile200 and Thr201 suggests that the T7 loop is not in a conformation that can make close contacts with Ala72.

**Table 5:** Best Fit Distances For Unique Peaks

| Peak (ppm) | Assignment | Best Fit Distance | T Model Distance | R1 Model Distance | R2 Model Distance | Region |
| --- | --- | --- | --- | --- | --- | --- |
| 14.9 | Ile200 Cδ1 | $4.9 \pm 0.4$ Å | 6.7 Å | 11.8 Å | 9.0 Å | T7 loop |
| 16.9 | Ile200 Cγ2 | $5.0 \pm 0.6$ Å | 6.6 Å | 11.3 Å | 7.8 Å | T7 loop |
| 17.1 | Ile176 Cγ2 | $3.5 \pm 0.6$ Å | 5.4 Å | 3.5 Å | 7.1 Å | Hinge |
| 18.9 | Ala72 Cβ | $5.0 \pm 0.7$ Å | 7.2 Å | 5.4 Å | 6.6 Å | Clasp |
| 43.1 | Leu168 Cβ | $4.07 \pm 0.03$ Å | 8.6 Å | 7.0 Å | 9.7 Å | Hinge |
| 46.7 | Gly71 Cα | $5.0 \pm 0.3$ Å | 4.1 Å | 7.9 Å | 9.1 Å | Clasp |
| 51.5 | Ala72 Cα | $5.2 \pm 0.2$ Å | 7.2 Å | 5.4 Å | 7.8 Å | Clasp |
| 64.6 | Thr201 Cα | $5.3 \pm 0.2$ Å | 4.7 Å | 11.7 Å | 11.5 Å | T7 loop |
| 66.4 | Thr215 Cα | $4.9 \pm 0.6$ Å | 6.9 Å | 13.5 Å | 6.8 Å | Clasp |
| 72.7 | Thr296 Cβ | $5.0 \pm 0.7$ Å | 8.1 Å | 7.5 Å | 11.5 Å | Hinge |

All fitting parameters can be found in table S3

**Figure 5:**
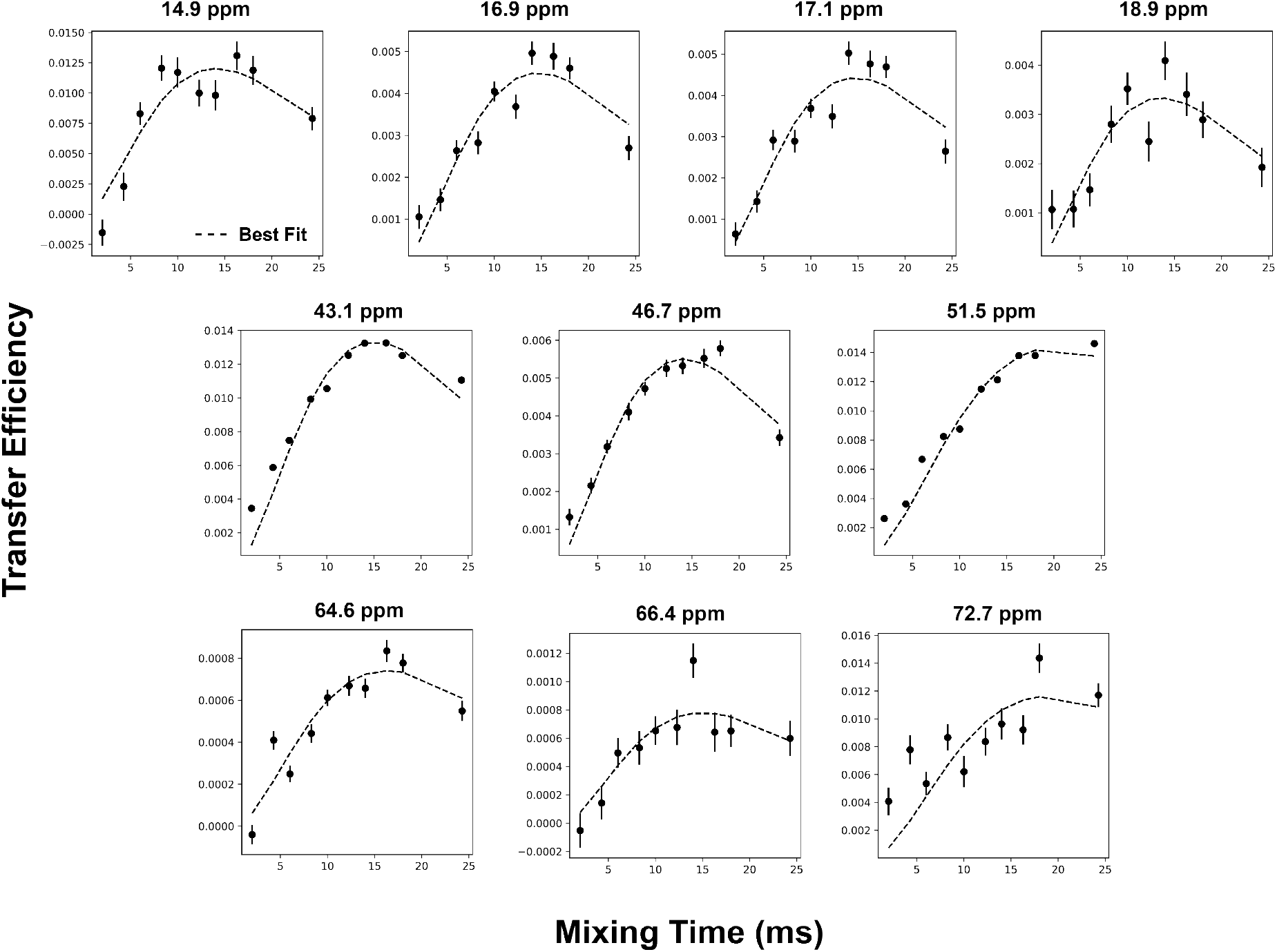
Best fit curves for ZF-TEDOR build-up curves for unique peaks in 2-glycerol and ^13^C*β*-Ala, ^13^C*α*-Gly sample spectra. Fits generated using SPINEVOLUTION and the python lmfit library using an isolated C–N spin pair model. Fits had 10 Hz J coupling applied to simulate higher order ^13^C–^13^C couplings present in the system.

Both Ile176 and Ala72 are located in flexible loops (figure 6), in the hinge and clasp regions respectively. Due to the flexible nature of these regions, we expect a degree of disagreement between our models and the actual protein structure. In the case of Ile176, it is possible that in the physiological interface, the sidechain of Arg271 faces the opposing monomer as it does in the R1 model. Additionally, given that the hinge region is common to all three interfaces, Ile176 is not expected to discriminate well between models. Ala72, however, is in the clasp region, which we expect to be highly predictive of interface state. The agreement between our measured distances and the predicted distances of Ala72 contacts however, must be read in the context of the other contacts in that region. Gly71 C*α* is predicted to have a contact distance of 4.1 Å in the T model, which agrees better with our measured distance of 5.0 ± 0.3 Åthan the R1 predicted distance of 7.9 Å. Thr215 is opposite of Ala72 in the T1 model, and the measured distance of 4.9± 0.6 Åfor Thr215 C*α* agrees better with the T1 prediction of 6.9 Å than the R1 prediction of 13.5 Å. Additionally, this suggests that the contacts between the H8 helix and the loop between H2 and H3 are closer in the actual interface than in the model.

**Figure 6:**
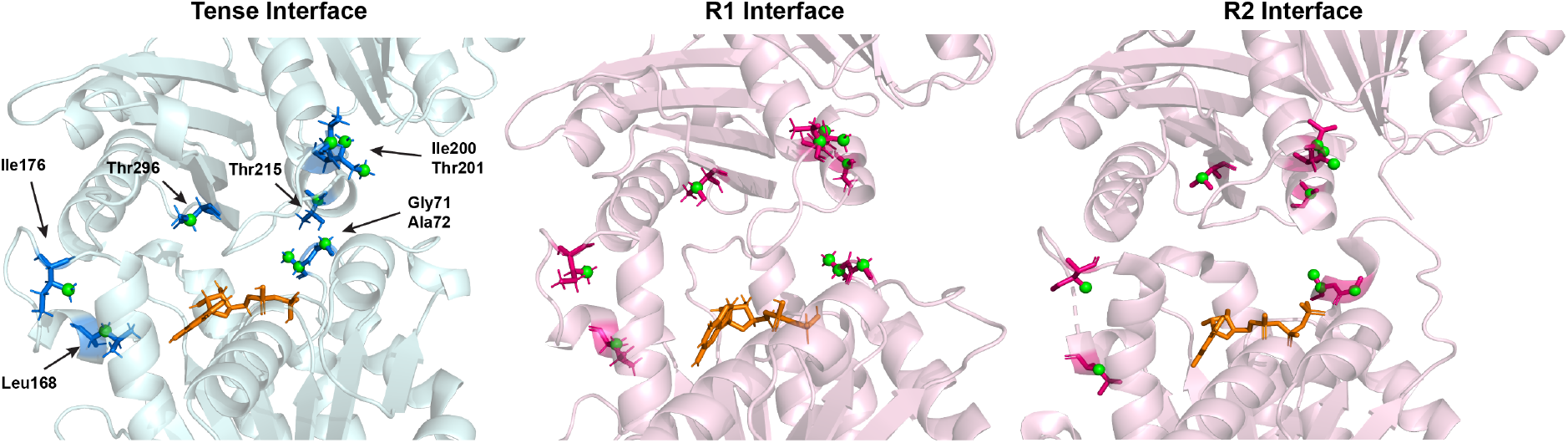
Residues with measured contacts in all three interface models with select residues labelled for comparison. Assigned atoms shown in green (spheres). See table 5 for measured and predicted contact distances for each model.

Overall, the distances we were able to measured agree with a T state interface. They also suggest that the interface is tighter than those observed in crystal structures. More work is needed to determine the precise contacts, which will allow for more accurate mapping of the interface. Taken together, these data support a model of the EcFtsZ filament where the monomers are in the T conformation. the interface itself is tight, with contacts as short of ~4.0 Å, and the T7 loop extends into the GTP-binding pocket. The clasp region of the interface makes several close contacts across the interface, indicating that it is involved in stabilizing the filament in a straight conformation.

## Conclusion

Using ZF-TEDOR experiments and DNP, we were able to directly observe contacts between ^13^C and ^15^N nuclei at the EcFtsZ inter-monomer interface. Once the ^13^C background in the ^15^N-labelled monomers was sufficiently depleted using ^12^C-glucose (99.9 %), inter-monomer contacts were observed. The lack of site-specific resonance assignments for the EcFtsZ monomer means that we cannot unambiguously assign peaks in the 1D ZF-TEDOR spectra. However, the availability of crystal structures of FtsZ monomers allowed us to generate chemical shift predictions.

We constructed homology models of the EcFtsZ dimer with tense (“T”) and relaxed (“R1”) monomers to model the monomer-monomer interface. We supplemented our homology models with a model constructed using an EcFtsZ R state monomer crystal structure (“R2”). Combining structural analysis of the models with chemical shift predictions allowed us to assign peaks in our ZF-TEDOR spectra. Using these tentative assignments, we demonstrated that the T model explains our data better than the R1 or R2 models. 13 individual residues were identified having at least one nucleus that corresponds to a unique peak. Of those, only one residue (Val171) corresponds to a peak that cannot be well explained by the T model. Two other residues (Ile200 and Thr201) correspond to peaks that can be explained by either the T model or the R1 model. The 10 other residues are best explained by the T model.

In addition to specific peak assignment, residue type analysis allowed us to distinguish between models. Specifically, the T model has six threonines present at the interface. Both R models have one only threonine present at the interface. We identified at least six separate peaks in the Thr C*α* region, all of which can be assigned to threonines predicted to be at the interface in the T model.

This study demonstrates that the wild type EcFtsZ filament consists mainly of T state monomers. We have also demonstrated that it is possible to use MAS NMR and DNP to study small, biologically relevant protein-protein interfaces in large, globular proteins. Further study is needed to refine the model and experimentally assign interface residues, but given the size of FtsZ compared to the interface, along with the inherent S/N limitations of TEDOR, this study demonstrates that significant information can be gained without full resonance assignments or multidimensional spectra.

## Acknowledgements

This work was supported by a grant from the National Science Foundation (NSF): MCB1412253 to A.E.M and by the National Institute of Health Grant P41 GM118302/GM/NIGMS NIH HHS/United States for the Center on Macromolecular Dynamics by NMR Spectroscopy located at the New York Structural Biology Center (NYSBC). A.E.M is a member of the NYSBC, and the data collected at NYSBC was enabled by a grant from NYSTAR and ORIP/NIH facility improvement grant CO6RR015495. The 600 MHz DNP/NMR spectrometer was purchased with funds from NIH grant S10RR029249. K.M.M was supported in part by a NSF Graduate Fellowship DGE 16-44869.

The authors thank Drs. Boris Itin, Keith Fritzsching, and Mike Goger of the NYSBC for help with DNP instrumentation and Dr. Ivan Sergeyev of Bruker BioSpin (Billerica, MA) for the use of their DNP instrument. K.M.M. also thanks Dr. Keith Fritzsching for his guidance and assistance and Kaitlin Entel and Kyra Cho for their work.

