## Supplemental Material for "GTP-bound *E. coli* FtsZ filaments are composed of Tense monomers: A DNP NMR study using interface detection"

March 31, 2022

### Additional Model Details

All three crystal structures used to build the models contain mutations to primary sequence that serve to stabilize the crystal packing. The Tense monomer (5MN4) is bound to GDP and has a mutation (F138A) that stabilizes the *S. aureus* FtsZ monomer in the Tense form. The SaFtsZ Relaxed monomer (5MN6) is also bound to GDP and has a stabilizing mutation (F136A) (*1*). Neither the T or R1 homology models constructed here explicitly contain the ligand, however the GTP-binding pockets of both homology models are structurally very similar to the template structures, with C$\alpha$ RMSDs of < 0.0001 Å, so it was not deemed necessary to dock a nucleotide into the models.

The EcFtsZ crystal structure used to construct the R2 model (6UNX) (*2*) is bound to GTP and contains the mutation L178E. Interestingly, this structure differs somewhat from the homology model used in the R1 model. The C$\alpha$ RMSD is 1.126 Å and while the differences are mainly in the mobile loop regions, the C-terminal subdomain of the EcFtsZ crystal structure is rotated relative to the homology model. The C$\alpha$ RMSD between the EcFtsZ crystal structure and tense homology model is 1.596 Å, suggesting that while the EcFtsZ crystal structure is definitely in a relaxed state, the orientation of the C-terminal subdomain is in-between that of the homology models. This may be a product of the specific mutation made to stabilize the EcFtsZ monomer, a crystal packing artifact, or variation between the species.

The R1 model forms a curved filament, the filament form observed after GDP hydrolysis. However, both the T model and the R2 model can form straight filaments. These filament forms are not equivalent however. The T model filament has a monomer spacing of 44.1 Å, which is in line with the canonical spacing of 44 Å (*1*). The R2 model filament has a monomer spacing of 46.8 Å, and, despite the increased spacing, an overall more compact filament.

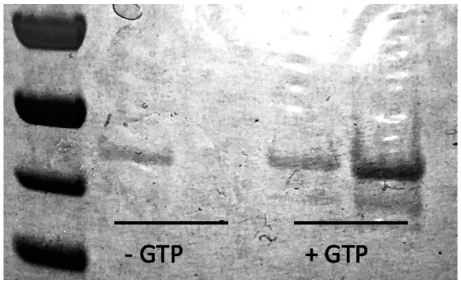


Figure 1: SDS-PAGE gel showing the results of a typical sedimentation assay with and without GTP added to FtsZ solution. Lanes: 1) Ladder. From top to bottom: 72 kDa, 55 kDa, 43 kDa, 34 kDa. 2) - GTP supernatant 3) - GTP pellet 4) + GTP supernatant 5) + GTP pellet.

### Biochemical Characterization of EcFtsZ

Purified EcFtsZ was assessed for functionality and polymerization dynamics using sedimentation, light scattering, and negative staining TEM.

Figure S1 shows the results of an EcFtsZ sedimentation assay with and without 1 mM GTP. 200 $\mu$g/mL EcFtsZ in a polymerization buffer (50 mM MES/NaOH pH 6.5, 10 mM $MgCl_2$, 50 mM KCl) was warmed to 30 °C and polymerization was initiated using 1 mM GTP (for the + GTP condition). Both samples were ultracentrifuged (Ti70.1 rotor, 70,000 rpm, 25 °C, 15 minutes). Samples from both the pellets and the supernatants were run on an SDS-PAGE gel and stained with Coomassie blue.

We also used 90° light scattering to compare buffer conditions in order to determine ideal conditions for MAS NMR samples. We were specifically interested in conditions that promoted polymer density and filament persistence. All light scattering experiments reported here were done on an ISS PC1 Spectrofluorometer at a wavelength of 350 nm. Light scattering experiments were done at 30 °C with stirring. 100 $\mu$L total volume was used. 500 $\mu$g/mL (12.5 $\mu$M) EcFtsZ was added to MES/NaOH buffer (pH 6.5) with 1 mM EGTA containing various concentrations of $MgCl_2$, KCl, and EDTA. 1 mM of nucleotide was used to induce polymerization. Scattering was observed for 60 minutes, or until scattering levels ceased to continue decreasing, indicating that depolymerization was complete.

Figure S2 demonstrates the typical light scattering pattern for EcFtsZ polymerization in a stan-

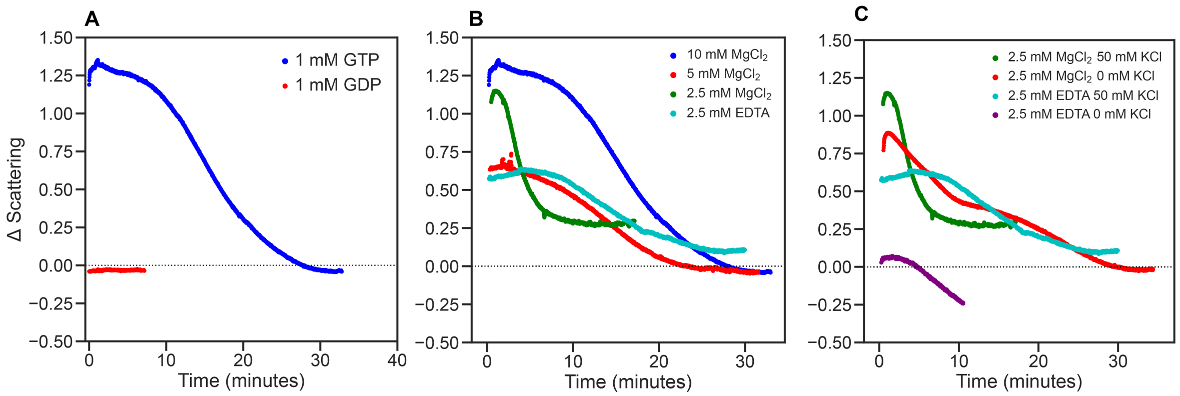


Figure 2: 90° light scattering demonstrating the effects of various buffer conditions on EcFtsZ polymerization. All assays were performed at 30 °C with stirring on. A) EcFtsZ polymerization induced by 1 mM GTP (blue). 1 mM GDP (red) does not induce any increase in scattering, demonstrating that GTP is necessary for polymerization. Buffer: 50 mM MES/NaOH (pH 6.5), 1 mM EGTA, 50 mM KCl, 10 mM $MgCl_2$. B) Effect of $MgCl_2$ concentration on EcFtsZ polymerization with 1 mM GTP. 10 mM $MgCl_2$ (blue) increases polymer density and filament duration greatly over 5 mM $MgCl_2$ (red) and 2.5 mM $MgCl_2$ (green). EcFtsZ can polymerize in an absence of $MgCl_2$ (cyan), which has a similar curve to 5 mM $MgCl_2$. C) 2.5 mM EDTA in the absence of $MgCl_2$ (teal) leads to polymerization and filament stabilization relative to 2.5 mM $MgCl_2$ (green). However, EDTA requires KCl in the buffer for polymerization (purple). EcFtsZ can polymerize in the absence of KCl (red), but the shape of the curve suggests aggregation.

dard polymerization buffer (50 mM MES/NaOH pH 6.5, 1 mM EGTA, 50 mM KCl, 10 mM $MgCl_2$ - MMK10). 1 mM GTP was used (blue trace) to initiate polymerization at time 0. The light scattering increased substantially within 20 seconds of addition of GTP (approximately the time it took at add GTP, mix, and restart scattering experiment). The scattering remained steady for approximately 8-10 minutes, indicating a steady state of polymerization and depolymerization, before returning to base-line over the course of the next 20 minutes. Figure S2 also demonstrations that the addition of 1 mM GDP (red) does not increase light scattering, indicating that, as expected, EcFtsZ does not polymerize in the presence of GDP. In fact, the scattering decreases slightly below the base-line when 1 mM GDP was added, suggesting that any transient oligomers dissociate back into monomer upon the addition of GDP, which is consistent with what has previously been reported (*3*)(*4*). Table S1 lists the various buffer conditions shown in Figure S2 along with the depolymerization half-time ($t_{1/2}$) in minutes. The half-time for each condition was calculated by normalizing each curve to its highest value and finding the time point nearest 0.5.

Table 1: EcFtsZ Polymerization Times

| Buffer | [$MgCl_2$] (mM) | [KCl] (mM) | [EDTA] (mM) | $t_{1/2}$ (min) |
|---|---|---|---|---|
| MMK10 | 10 | 50 | 0 | 15.2 |
| MMK5 | 5 | 50 | 0 | 12.2 |
| MMK2.5 | 2.5 | 50 | 0 | 4.32 |
| MM | 2.5 | 0 | 0 | 9.67 |
| MEK | 0 | 50 | 2.5 | 15.58 |
| ME | 0 | 0 | 2.5 | N/A |

Reducing the concentration of (mM) from 10 mM to 2.5 mM greatly decreased the polymerization half-time (figure S2; table S1), suggesting both a lower polymer density and faster GTP hydrolysis at lower $MgCl_2$ concentrations. 5 mM $MgCl_2$ has a similar curve shape as 10 mM $MgCl_2$ and a half-time that is only slightly shorter, but seems to have a smaller polymer density based on the overall high of the curve in terms of Δ scattering. However, in the absence of $MgCl_2$, when residual cations are removed with 2.5 mM EDTA, EcFtsZ polymerizes and has a half-time and a curve comparable to the 10 mM $MgCl_2$ condition (figure S2C; table S1). This is consistent with previous findings in the literature that $Mg^{2+}$ is required for FtsZ's GTPase activity but not for polymerization (*3*)(*5*). However, in the presence of EDTA and KCl (MEK) the total Δ scattering (the difference between the base-line scattering and largest scattering upon addition of GTP) is less than half as much as the Δ scattering in the case of 10 mM $MgCl_2$ (table S1). While the total amount of scattering is not a quantitative measure of polymer density, and, in fact, varies between replicates (figure S2C; table S1), this large of a difference does suggest that the polymer density generated using the EDTA/KCl buffer is less than for the $MgCl_2$ buffers.

Mukherjee *et al.* (1999) reported that EcFtsZ's GTPase activity is greatly reduced in the absence of KCl, even when in the presence of $MgCl_2$ (*5*). We tested this condition, and found that EcFtsZ does, indeed, polymerize in the absence of KCl (figure S2C; table S1 - MM). However, the shape of the depolymerization curve is substantially different than those for the other polymerization conditions, suggesting that depolymerization dynamics differ. We also tested the EDTA condition in the absence of KCl (ME) and found that EcFtsZ did not polymerize in any substantial way (figure S2C), verifying that EcFtsZ does need KCl when $MgCl_2$ is not present. Additionally, we found that EcFtsZ tends to form insoluble aggregates when concentrated in buffers without KCl (data not shown). Taken together, these data suggest that moderate concentrations of KCl (25–50 mM) is necessary for EcFtsZ to behave well. This finding supported my inclusion of KCl in all EcFtsZ NMR and storage buffers.

**Negative Staining EM**

We used negative staining TEM to visualize filament morphology under various buffer conditions. All EM images were collected at the New York Structural Biology Center (NYSBC) Simon's Electron Microscopy Center with assistance from their staff. All negative staining TEM images were collected on a JEOL JEM-1230 microscope with a Gatan US-4000 CCD detector 4k x 4k at the NYSBC. Purified EcFtsZ was diluted to 0.625 mg/mL (3.125 $\mu$M) in the appropriate buffer (50 mM MES/NaOH pH 6.5, 1 mM EGTA base plus metal cations as specified in Table 3.1) and incubated at 30 °C for 5–10 minutes. 1 mM GTP was then added to induce polymerization and the sample was incubated again at 30 °C for 5 minutes. 3 $\mu$L of protein solution was spotted onto a plasma cleaned continuous carbon grid. After 30–60 seconds the grid was blotted dry and washed two times with 20 $\mu$L buffer and three times with 20 $\mu$L 1 % uranyl formate stain (final staining for 1 minute).

EcFtsZ polymerized in MMK10 buffer (10 mM $MgCl_2$, 50 mM KCl) had very high polymer density and did not yield good images. EcFtsZ polymerized in MMK2.5 buffer (2.5 mM $MgCl_2$, 50 mM KCl; figure S3) yielded images with densely packed filaments in both straight and curved conformations. When EcFtsZ was polymerized in MMK2.5 buffer with 1 mM GTP$\gamma$S rather than GTP, no filaments were observed (figure S3), which is in line with previous reports that GTP$\gamma$S does not promote FtsZ polymerization (*6*). We also took TEM images of two other polymerizing conditions (buffers MM and MEK; see table S1) and confirmed that when EcFtsZ is polymerized without KCl (buffer MM, 2.5 mM $MgCl_2$; figure S3) there are some filaments present, but much fewer than in the presence of 50 mM KCl, and the morphology of the filaments present seems to

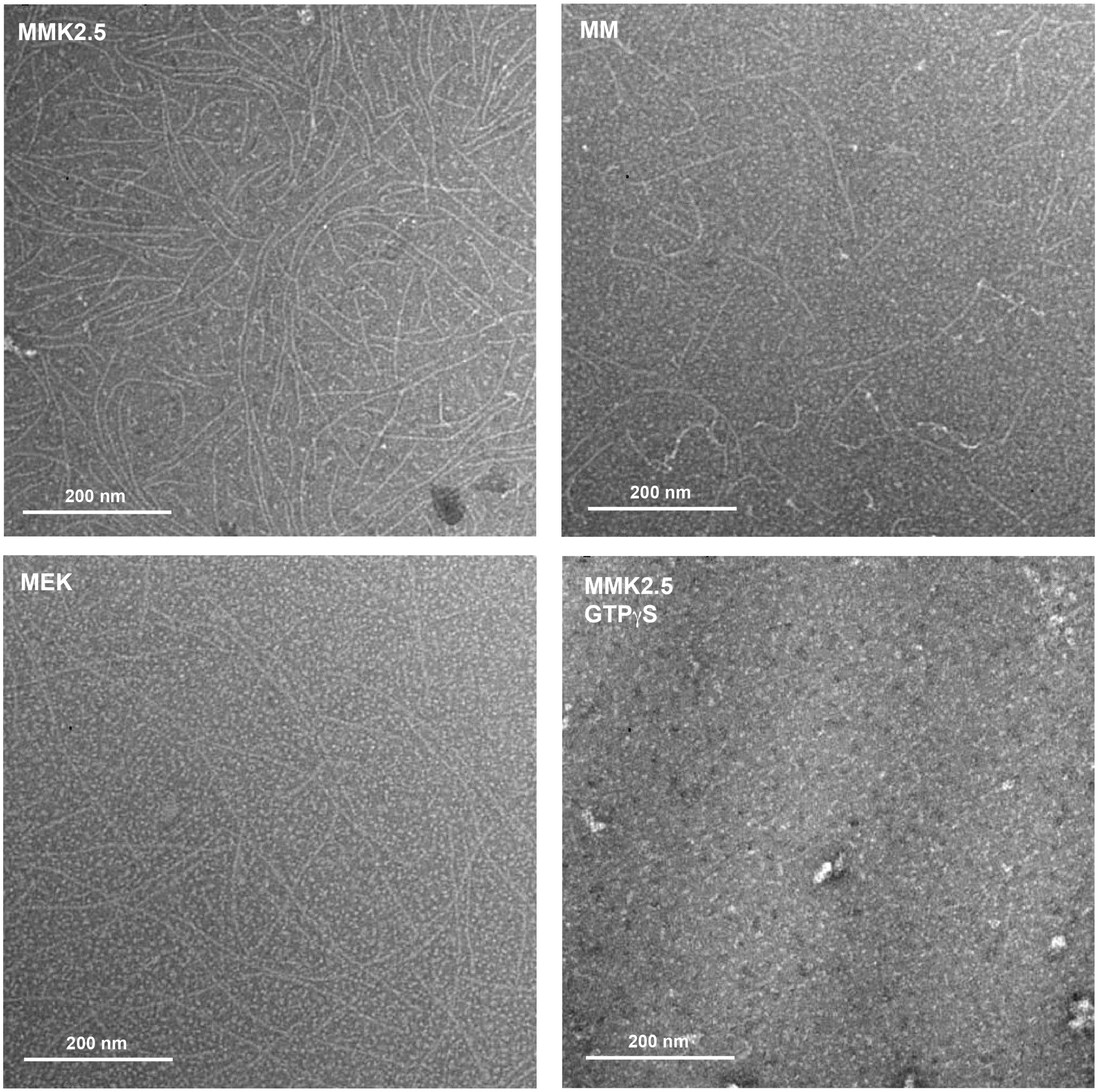


Figure 3: Representative TEM images of EcFtsZ filaments in various buffer conditions. MMK2.5: 50 mM KCl, 2.5 mM $MgCl_2$ MM: 2.5 mM $MgCl_2$ MEK: 50 mM KCl, 2.5 mM EDTA MMK2.5 + GTP$\gamma$S: EcFtsZ polymerized in MMK2.5 buffer with 1 mM GTP$\gamma$S instead of 1 mM GTP.

be less well behaved. In the presence of 50 mM KCl but not $Mg^{2+}$ (buffer MEK, 2.5 mM EDTA; figure S3), the majority of the filaments are straight, and the persistence length seems to be longer than in the MMK2.5 case. However, the polymer density is lower and there is a higher degree of monomers present in the background.

These TEM negative staining images confirm our light scattering data, showing that both KCl and $MgCl_2$ are necessary for the highest degree of polymer density. Polymerizing in the presence of EDTA to chelate out any residual $Mg^{2+}$ promotes the formation of relatively stable, predominantly straight filaments, but these filaments do depolymerize over the course of 40 min to an hour and polymer density is an issue. These findings motivated our use of MES buffer (pH 6.5) with 50 mM KCl and 1 mM EGTA for all NMR experiments described in this study.

### DNP Spectra of GTP-bound EcFtsZ Filaments

Figure S4 shows the $^{13}$C Hartmann-Hahn cross polarization (CP) spectrum for each sample used in this study, including the $^{13}$C CP spectrum of the U-$^{15}$N control sample showing the spectrum of the natural abundance $^{13}$C present in the U-$^{15}$N enriched monomers. The TEDOR build-up curves for each sample were normalized to their respective $^{13}$C CP spectrum. The differences in the $^{13}$C CP spectra illustrates the differences in the $^{13}$C labelling schemes used in each sample. The $^{13}$Ca-Gly, $^{13}$Cb-Ala spectrum shows significant isotopic scrambling that is discussed in detail below.

### Isotopic Labelling Pattern

The $^{13}$C$\beta$ of alanine scrambled to the methyl carbons of leucine and valine, along with the C$\gamma$2 carbon of isoleucine, which is consistent with the alanine metabolic pathways (*7*)(*8*). The $^{13}$C$\alpha$ of glycine scrambled to the $^{13}$C$\alpha$ of both serine and threonine, which again is consistent with known metabolic pathways (*8*). Additionally, the CP spectra showed peaks in the isoleucine $^{13}$C$\delta$ range, as well as peaks consistent with leucine and valine $^{13}$C$\alpha$s. These can be explained by the addition of natural abundance isoleucine and $\alpha$-ketoisovalerate. U-$^{12}$C glucose (99.9 %) was used as the carbon source for the sample, but the added isoleucine and $\alpha$-ketoisovalerate had natural abundance $^{13}$C levels of 1.1 %. This means that all the carbons in isoleucine, leucine, and valine have $^{13}$C levels of 1.1 %, while all other non-labelled residues have $^{13}$C levels of 0.1 %. Therefore, unlabelled ILV carbons should be enriched in $^{13}$C relative to the rest of the unlabelled residues, leading to some visible peaks in the spectrum. Figure S5 shows the scrambling pattern present in

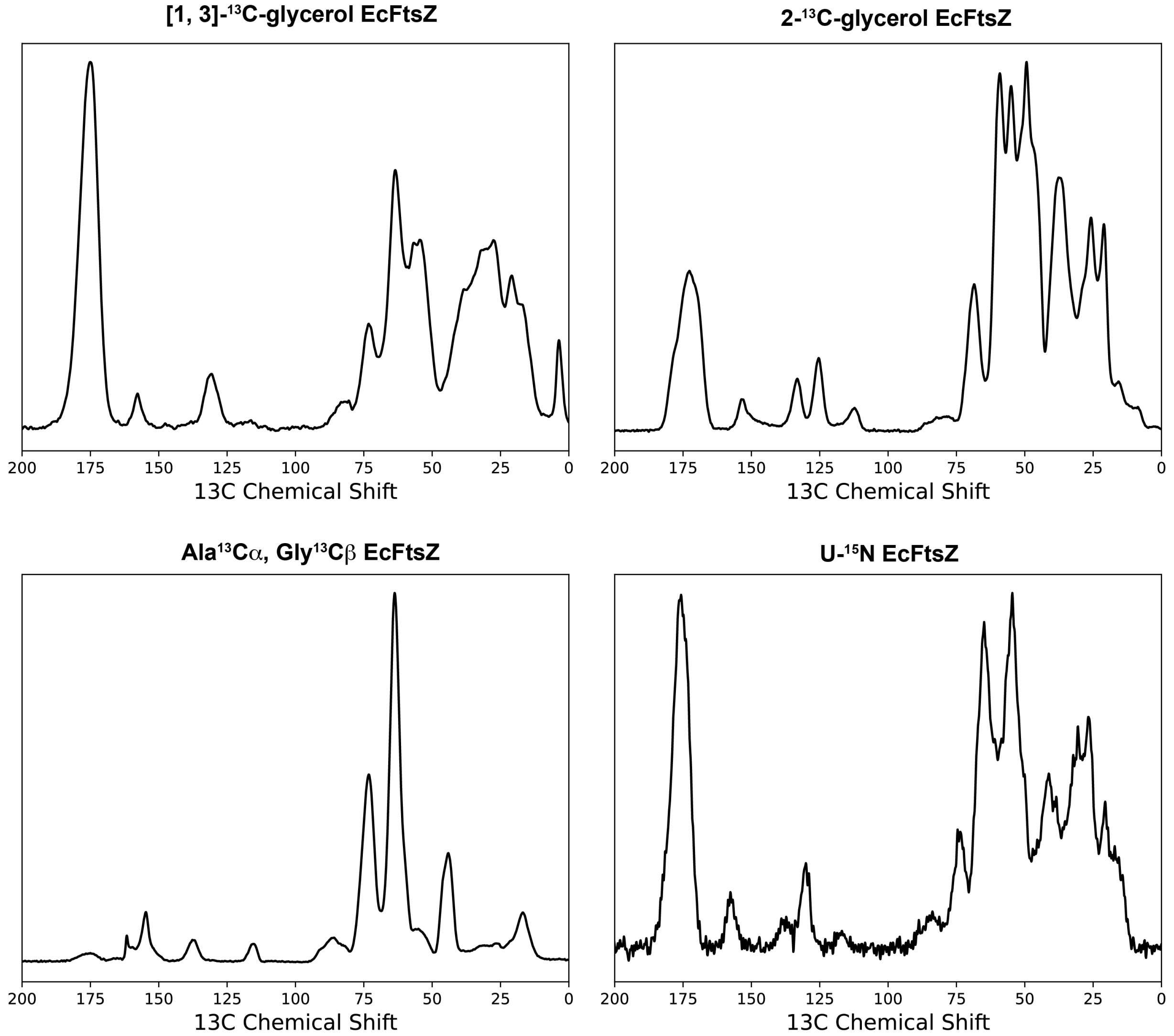


Figure 4: $^{13}$C Cross Polarization spectra for DNP samples used in this study. The parameters for each spectrum are as follows: 1, 3-glycerol: ns = 32 scans, $^{1}$H 90 ° pulse of 172 kHz, $^{13}$C CP power centered at 61 kHz with a fixed proton power of 77 kHz, and a proton decoupling strength of 82 kHz. 2-glycerol: ns = 16 scans, $^{1}$H 90 ° pulse of 208 kHz, $^{13}$C CP power centered at 70 kHz with a fixed proton power of 84 kHz, and a proton decoupling strength of 83 kHz. $^{13}$Ca-Gly, $^{13}$Cb-Ala: ns = 64 scans, $^{1}$H 90 ° pulse of 160 kHz, $^{13}$C CP power centered at 63 kHz with a fixed proton power of 77 kHz, and a proton decoupling strength of 83 kHz. U- $^{15}$N: ns = 64 scans, $^{1}$H 90 ° pulse of 169 kHz, $^{13}$C CP power centered at 63 kHz with a fixed proton power of 77 kHz, and a proton decoupling strength of 83 kHz. All samples had an MAS frequency of 14,000 Hz. All spectra were processed with 50 Hz of exponential line broadening.

the spectrum.

A complicating factor of this analysis is that the solvent is 30 % glycerol. Glycerol has two $^{13}$C peaks—one at 74.87 ppm corresponding to carbon 2 and one at 65.32 ppm corresponding to the

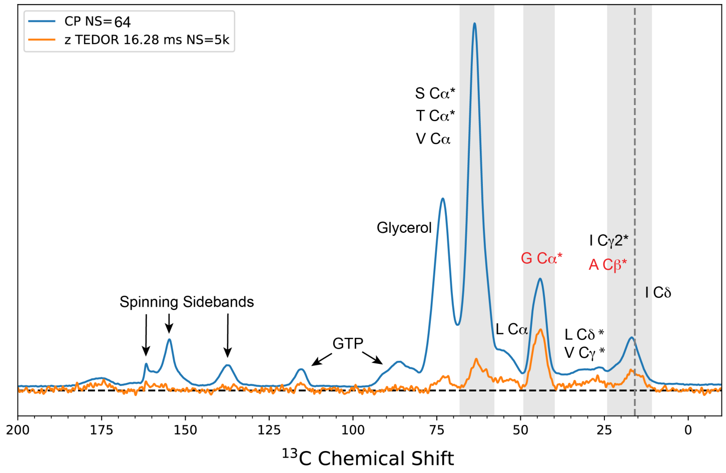


Figure 5: 1D $^{13}$C DNP spectra of of $^{13}$C$\beta$-Ala, $^{13}$C$\alpha$-Gly EcFtsZ mixed monomer sample showing CP (blue, ns=64) and a representative ZF-TEDOR spectrum (orange, 16.28 ms mixing time, ns=5120), annotated to show the scrambling pattern.

degenerate 1 and 3 carbons. 65.32 ppm is, unfortunately, smack dab in the middle of the threonine C$\alpha$ region. Given that ~8.5 % of the solvent accessible surface of EcFtsZ corresponds to nitrogens, this presents a potential problem. The samples were prepared using U-$^{12}$C, $^{2}$H glycerol, which is 99.95 % $^{12}$C. As discussed in above, even 0.1 % $^{13}$C can be seen in the DNP spectra.

None of the peaks in this region of the spectra listed in table 4 are within ± 0.5 ppm of 65.3 ppm, suggesting that, at least by the criteria used to identify peaks in this study, they are not glycerol peaks. When compared to a spectrum where the glycerol peaks are apparent—a natural abundance $^{13}$C EcFtsZ DNP spectrum collected as a part of the $^{15}$N control sample data set—the peaks present in the $^{13}$C$\beta$-Ala, $^{13}$C$\alpha$-Gly spectra are not centered at the same positions (figure S6). While there are likely contributions from glycerol in the threonine C$\alpha$ region, any peaks not within 0.5 ppm of 65.3 ppm can be reasonably assumed to be protein peaks.

Figure S6 shows a comparison between the 1D $^{13}$C$\beta$-Ala, $^{13}$C$\alpha$-Gly $^{13}$C CP spectrum and the 1D natural abundance $^{13}$C CP spectrum (acquired using the U-$^{15}$N FtsZ sample). The glycerol peaks are apparent in the n.a. spectrum, centered at 65 ppm and 74 ppm. However a clear shift is seen in AG sample spectrum in these regions (the peaks are centered at 63 ppm and 72 ppm

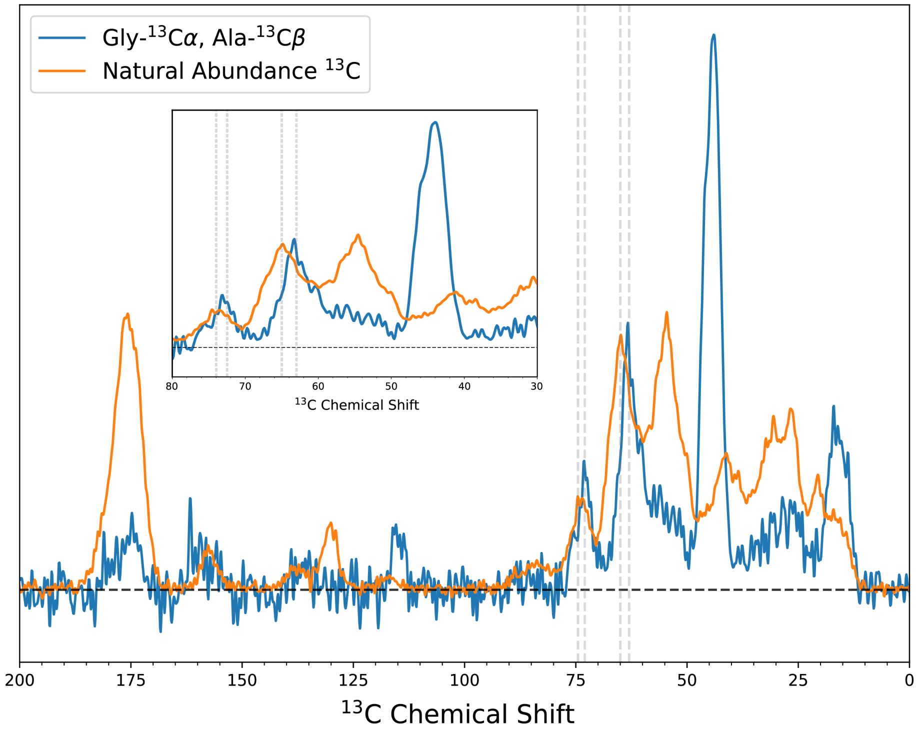


Figure 6: $^{13}$Cβ-Ala, $^{13}$Cα-Gly $^{13}$C CP spectrum (blue) and natural abundance $^{13}$C EcFtsZ CP DNP spectrum (orange).

respectively). This demonstrates that the peaks in these regions in the AG spectrum, analyzed in this study, are not solely due to the presence of glycerol in the sample.

### Selected ZF-TEDOR Build-up Curves

Figures S7 and S8 shows the build-up curves for the identified unique peaks in this study, across all three samples. Curves were plotted by integrating across a total of 1 ppm centered on the peak and normalizing the transfer efficiency as compared to the $^{13}$C-CP spectrum for each respective sample. The curves for the same regions of the U-$^{15}$N control sample are shown as a control. These curves clearly demonstrate that inter-monomer transfer is occurring and that polarization build-up peaks between 14 ms and 20 ms in most cases.

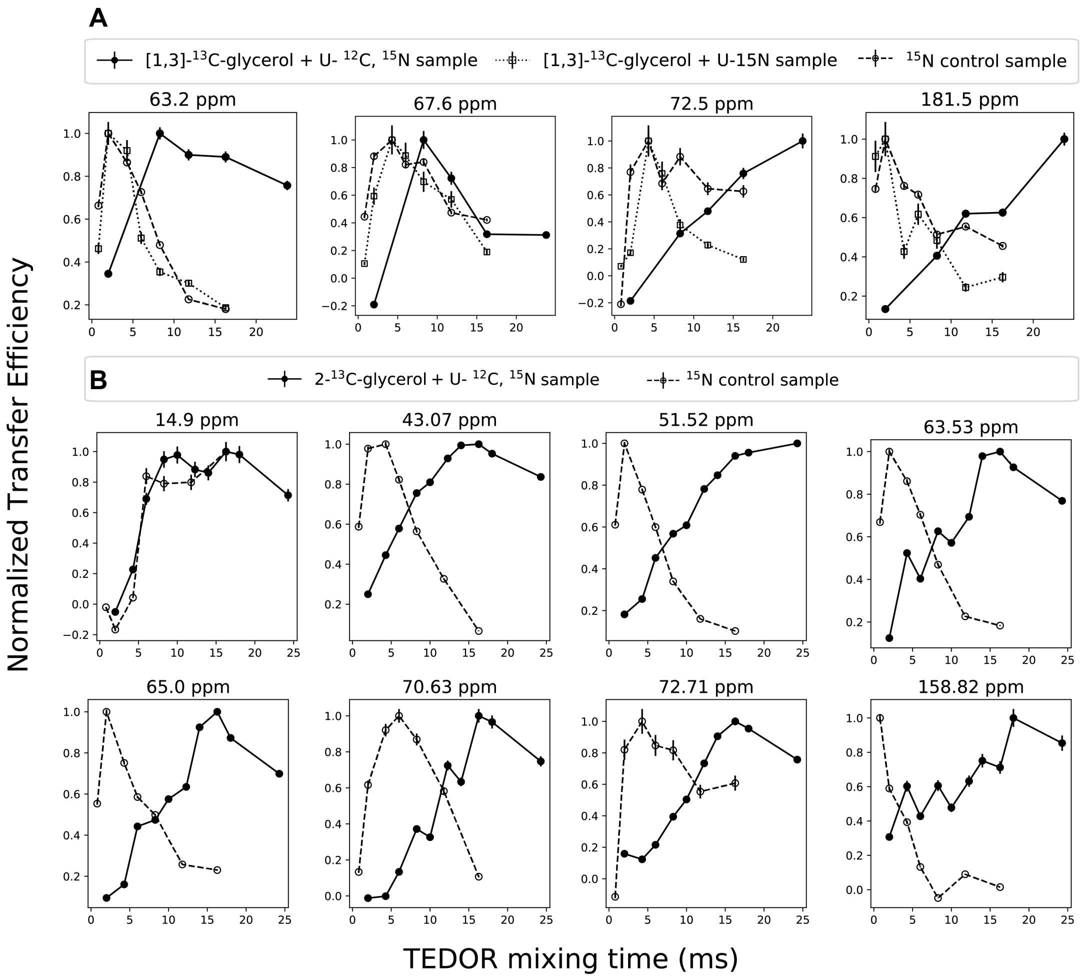


Figure 7: A) ZF-TEDOR build-up curves for the [1,3]-$^{13}$C-glycerol + U-$^{15}$N, $^{12}$C EcFtsZ mixed label sample (black, •), [1,3]-$^{13}$C-glycerol + U-$^{15}$N EcFtsZ mixed label sample (dotted, □), and U-$^{15}$N EcFtsZ sample (dashed, ◦). B) ZF-TEDOR build-up curves for the 2-$^{13}$C-glycerol + U-$^{15}$N, $^{12}$C EcFtsZ mixed label sample (black, •) and the U-$^{15}$N EcFtsZ sample (dashed, ◦). All peaks were integrated across a total of 1 ppm centered on the peak and plotted as the normalized transfer efficiency compared to the $^{13}$C-CP spectrum for each respective sample.

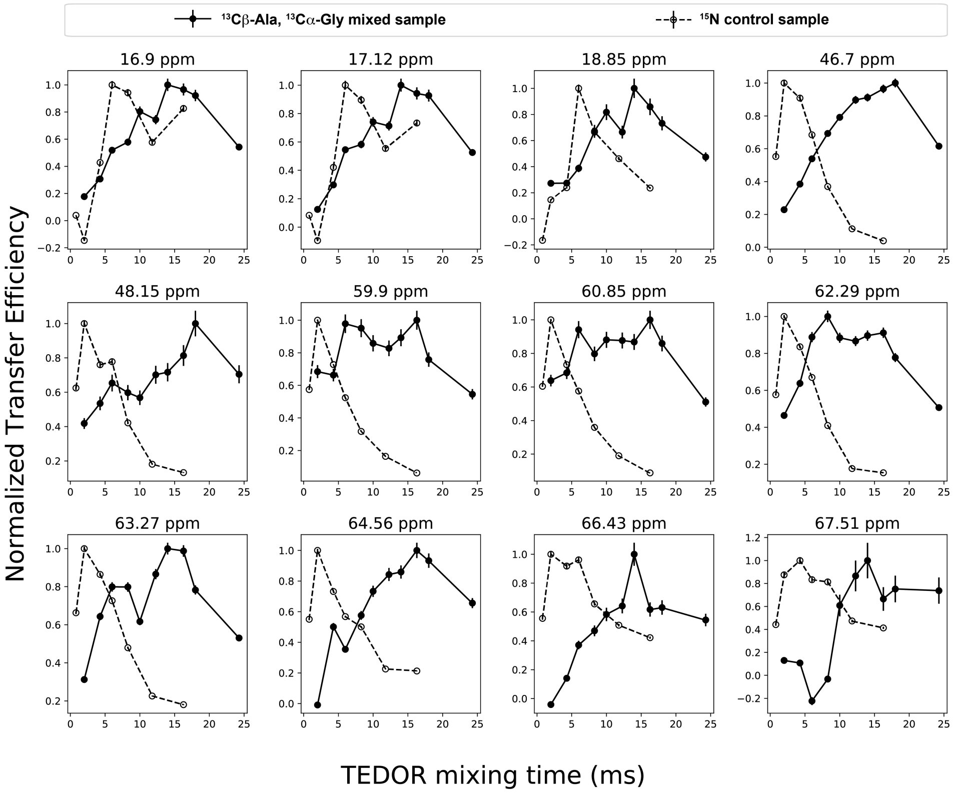


Figure 8: A) ZF-TEDOR build-up curves for the $^{13}$C$\beta$-Ala, $^{13}$C$\alpha$-Gly + U-$^{15}$N, $^{12}$C EcFtsZ mixed label sample (black, •) and U-$^{15}$N EcFtsZ sample (dashed, ∘). All peaks were integrated across a total of 1 ppm centered on the peak and plotted as the normalized transfer efficiency compared to the $^{13}$C-CP spectrum for each respective sample.

Table 2: Unique Peaks in the ZF-TEDOR Spectra

| Peak (ppm) | Predicted Assignment | Predicted Chemical Shift (ppm) | Sample |
|---|---|---|---|
| 14.9 | Ile200 C$\delta$1 | 14.51 | 2-glycerol |
| 16.9 | Ile200 C$\gamma$2 | 17.00 | $^{13}$C$\beta$-Ala, $^{13}$C$\alpha$-Gly |
| 17.1 | Ile176 C$\gamma$2 | 17.55 | $^{13}$C$\beta$-Ala, $^{13}$C$\alpha$-Gly |
| 18.9 | Ala72 C$\beta$ | 19.17 | $^{13}$C$\beta$-Ala, $^{13}$C$\alpha$-Gly |
| 43.1 | Leu168 C$\beta$ | 43.11 | 2-glycerol |
| 46.7 | Gly71 C$\alpha$ | 46.34 | $^{13}$C$\beta$-Ala, $^{13}$C$\alpha$-Gly |
| 48.2 | Gly20 C$\alpha$ | 47.78 | $^{13}$C$\beta$-Ala, $^{13}$C$\alpha$-Gly |
| 51.5 | Ala72 C$\alpha$ | 51.75 | 2-glycerol |
| 59.9 | Thr296 C$\alpha$ | 59.98 | $^{13}$C$\beta$-Ala, $^{13}$C$\alpha$-Gly |
| 60.9 | Ser177 C$\alpha$ | 61.16 | $^{13}$C$\beta$-Ala, $^{13}$C$\alpha$-Gly |
| 62.2 | Thr291 C$\alpha$ | 62.32 | $^{13}$C$\beta$-Ala, $^{13}$C$\alpha$-Gly |
| 63.3 | Thr65 C$\alpha$ | 62.85 | $^{13}$C$\beta$-Ala, $^{13}$C$\alpha$-Gly |
| 64.6 | Thr201 C$\alpha$ | 64.46 | $^{13}$C$\beta$-Ala, $^{13}$C$\alpha$-Gly |
| 65.0 | Ile200 C$\alpha$ | 65.13 | 2-glycerol |
| 66.4 | Thr215 C$\alpha$ | 66.12 | $^{13}$C$\beta$-Ala, $^{13}$C$\alpha$-Gly |
| 67.5 | Thr281 C$\alpha$ | 67.11 | $^{13}$C$\beta$-Ala, $^{13}$C$\alpha$-Gly |
| 70.6 | Thr65 C$\beta$ | 70.53 | 2-glycerol |
| 72.7 | Thr296 C$\beta$ | 72.59 | 2-glycerol |
| 158.8 | Arg202 Cz | 159.27 | 2-glycerol |

Table 3: Best Fit Parameters

| Peak (ppm) | Predicted Assignment | Best Fit Distance (Å) | $t_2$ (ms) | Fit RMSD |
|---|---|---|---|---|
| 14.9 | Ile200 C$\delta$1 | 5.0 ± 0.4 | 7.3 | 0.41 |
| 16.9 | Ile200 C$\gamma$2 | 5.0 ± 0.6 | 7.7 | 0.35 |
| 17.1 | Ile176 C$\gamma$2 | 3.5 ± 0.6 | 11.7 | 0.34 |
| 18.9 | Ala72 C$\beta$ | 5.0 ± 0.7 | 7.0 | 0.49 |
| 43.1 | Leu168 C$\beta$ | 4.07 ± 0.03 | 9.0 | 0.33 |
| 46.7 | Gly71 C$\alpha$ | 5.0 ± 0.3 | 7.3 | 0.25 |
| 48.2 | Gly20 C$\alpha$ | 5.7 ± 0.4 | 7.6 | 1.66 |
| 51.5 | Ala72 C$\alpha$ | 5.2 ± 0.1 | 10.8 | 0.23 |
| 59.9 | Thr296 C$\alpha$ | 5.00 ± 0.01 | 5.3 | 1.40 |
| 60.9 | Ser177 C$\alpha$ | 5.02 ± 0.01 | 5.6 | 1.35 |
| 62.2 | Thr291 C$\alpha$ | 5.05 ± 0.01 | 5.3 | 0.69 |
| 63.3 | Thr65 C$\alpha$ | 5.2 ± 0.5 | 6.2 | 0.79 |
| 64.6 | Thr201 C$\alpha$ | 5.19 ± 0.07 | 9.0 | 0.37 |
| 65.0 | Ile200 C$\alpha$ | 4.7 ± 0.6 | 10.1 | 0.34 |
| 66.4 | Thr215 C$\alpha$ | 4.9 ± 0.6 | 7.9 | 0.48 |
| 67.5 | Thr281 C$\alpha$ | 5.2 ± 0.7 | 13.69 | 0.73 |
| 70.6 | Thr65 C$\beta$ | 5.0 ± 1.0 | 14.7 | 0.43 |
| 72.7 | Thr296 C$\beta$ | 5.0 ± 0.7 | 11.9 | 0.31 |
| 158.8 | Arg202 Cz | 4.14 ± 0.07 | 12.3 | 0.89 |
